# Loss of *polr1c* disrupts myelination in a zebrafish model of *POLR1C*-associated disease

**DOI:** 10.64898/2026.09.24.752997

**Authors:** Lauren B. Sands, Camille E.A. Goo, Laura K. White, Bailey T. Lubash, M. Cathleen McKinney, Fengli Guo, Chris W. Seidel, Xia Zhao, Paul A. Trainor, Kristin E.N. Watt

**Affiliations:** Department of Craniofacial, Oral and Materials Sciences, University of Colorado Anschutz, Aurora, CO, USA; Department of Biochemistry & Molecular Genetics, University of Colorado Anschutz, Aurora, CO, USA; Stowers Institute for Medical Research, Kansas City, MO, USA; Department of Pediatrics, McGovern School of Medicine, UTHealth Houston, TX, USA

## Abstract

Pathogenic variants in *POLR1C*, which encodes a shared subunit of RNA Polymerases (Pols) I and III, cause Treacher Collins syndrome (TCS) and POLR3-related leukodystrophy. While Pol I and Tp53-dependent mechanisms have been implicated in the pathogenesis of TCS, the basis of hypomyelination in *POLR1C*-associated POLR3-related leukodystrophy remains incompletely understood. Here, we show that *polr1c* mutant zebrafish exhibit reduced myelination in addition to previously described craniofacial anomalies. Oligodendrocyte precursor cells exhibit increased activation of the Tp53 pathway; however, these cells do not undergo apoptosis. Consistent with this finding, *tp53* inhibition reduces cell death in *polr1c* mutants but fails to restore myelination, indicating that myelination deficits are not driven by Tp53-dependent progenitor loss in this model. *polr1c* mutants also exhibit reduced rRNA transcription by Pol I and reduced expression of some Pol III-transcribed tRNAs. Altogether, these data indicate distinct tissue-specific responses to *polr1c* deficiency and suggest persistent impairment of rRNA transcription contributes to deficient myelin development. These findings expand the developmental consequences of *polr1c* loss and advance our understanding of the molecular basis of *POLR1C*-associated diseases.

## Introduction

Ribosome biogenesis is an essential and ubiquitous process requiring the coordination of RNA Polymerases (Pol) I, II, and III to produce ribosomal RNA (rRNA), ribosomal proteins, and associated assembly factors (Lafontaine and Tollervey, 2001). This process is highly regulated to coordinate ribosome production with nutrient availability, cell cycle progression, and cellular demands (Grummt, 2003; Moir and Willis, 2013; Ni and Buszczak, 2023). RNA Polymerase I (Pol I) transcribes the 18S, 5.8S, and 28S rRNAs, which is a rate-limiting step in ribosome biogenesis (Laferté et al., 2006). Pol III transcribes 5S rRNA, transfer RNAs (tRNAs), and a variety of additional noncoding RNAs (Girbig et al., 2021; Ramsay et al., 2020; White, 2011). Ribosome biogenesis is dynamically required during development, and influences proliferation, cell fate commitment, and differentiation (Woolnough et al., 2016; Zhang et al., 2014). Despite the essential and ubiquitous nature of ribosome biogenesis, disruptions in this process lead to a group of disorders termed ribosomopathies which display distinct, tissue-specific phenotypes (Watt et al., 2023; Yelick and Trainor, 2015).

POLR1C and POLR1D are subunits shared by Pol I and Pol III that contribute to polymerase assembly and function (Boguta, 2022; Wild and Cramer, 2012). Pathogenic variants in *POLR1C* are associated with two distinct syndromes and are typically inherited in an autosomal recessive pattern. Variants associated with Treacher Collins syndrome (TCS 3; OMIM 248390) produce predominantly craniofacial differences, including hypoplasia of craniofacial bones such as the mandible, cleft palate, and external ear anomalies (Dauwerse et al., 2011). Other *POLR1C* variants are associated with *POLR3*-related leukodystrophy (HLD11; OMIM 616494), a neurological disorder characterized by hypomyelination in the central nervous system (Gauquelin et al., 2019; Thiffault et al., 2015). How disruption of a shared Pol I and III subunit produces these distinct tissue-specific phenotypes remains unresolved.

Our prior work demonstrated a requirement for rRNA transcription by Pol I during neural crest cell and craniofacial development. Neural crest cells (NCCs) are a migratory, progenitor population which give rise to much of the craniofacial cartilage and bone (Bhatt et al., 2013) and are particularly sensitive to impaired ribosome biogenesis (Falcon et al., 2022; Jones et al., 2008; Noack Watt et al., 2016; Weaver et al., 2015). NCC-specific conditional deletion of *Polr1c* in mice results in NCC death (Falcon et al., 2022), while mutations in *polr1c* or *polr1d* in zebrafish result in cranioskeletal defects, consistent with features of Treacher Collins syndrome (TCS) (Noack Watt et al., 2016). These cranioskeletal defects are associated with reduced rRNA transcription by Pol I and Tp53-dependent neuroepithelial cell death (Noack Watt et al., 2016). Altogether, these studies confirmed that NCC progenitor loss, as a consequence of impaired ribosome biogenesis, is a conserved mechanism underlying the craniofacial differences associated with *Polr1c* deficiency (Falcon et al., 2022; Noack Watt et al., 2016). Whether a similar mechanism accounts for the neurological phenotypes associated with *POLR1C* remains unknown.

Ribosome biogenesis is dynamically regulated during neurogenesis (Chau et al., 2018), and impaired rRNA transcription can compromise the survival of neural progenitors and neurons (Parlato et al., 2008; Smallwood et al., 2023). Dynamic requirements may also occur in myelinating cells, as myelin in the central nervous system (CNS) is enriched for transcripts associated with translation and ribosome assembly, and disruption of the ribosome biogenesis factor *pescadillo* impairs myelin gene expression in zebrafish (Simmons and Appel, 2012; Thakurela et al., 2016). Understanding these requirements is relevant to *POLR1C*-associated HLD11, in which deficient CNS myelination is a defining feature. Myelin is produced by oligodendrocytes in the CNS, which arise from oligodendrocyte precursor cells (OPCs). In the developing spinal cord, the motor neuron progenitor domain (pMN) first produces motor neurons and then gives rise to OPCs, which delaminate, proliferate, and migrate towards their target axons to begin myelination (Ackerman and Monk, 2016; Xia and Fancy, 2021). In the peripheral nervous system (PNS), myelin is instead produced by NCC-derived Schwann cells (Ackerman and Monk, 2016). How *POLR1C* deficiency affects the development and function of myelinating cells remains unclear.

One proposed explanation for the distinct *POLR1C-*assocated syndromes is that TCS-associated variants preferentially disrupt Pol I while HLD11-associated variants predominantly impair Pol III (Misiaszek et al., 2021; Thiffault et al., 2015). However, additional studies indicate that TCS variants may affect both Pol I and Pol III (Ramsay et al., 2020; Walker-Kopp et al., 2017), and individuals with *POLR1C* variants can exhibit both craniofacial and neurological phenotypes (Gauquelin et al., 2019; Mirchi et al., 2023). Together, this suggests that there may not be a strict correspondence between the clinical phenotype and dysfunction of a single polymerase, and raises the possibility that both Pol I and Pol III may contribute to the phenotypic spectrum observed in *POLR1C*-associated diseases.

We therefore used a *polr1c* loss-of-function zebrafish model to determine how global *polr1c* deficiency affected myelination and if the mechanisms underlying neurological phenotypes are consistent with those previously defined for craniofacial development (Noack Watt et al., 2016). Here, we show that *polr1c^-/-^* zebrafish exhibit impaired myelin development, increased Tp53 signaling, and tissue specific apoptosis. Genetic inhibition of *tp53* reduces cell death but does not restore myelination in *polr1c^-/-^* zebrafish, indicating that the neurological phenotype cannot be explained by Tp53-dependent progenitor loss. Molecular analyses further reveal a significant reduction in rRNA transcription by Pol I, while only modest differences were observed in tRNAs transcribed by Pol III. These data reveal distinct tissue-specific responses to the loss of *polr1c* and suggest that impaired rRNA transcription and ribosome biogenesis contribute to impaired myelination in this model.

## Results

### Neuronal development and myelination are disrupted in *polr1c^-/-^* zebrafish

Craniofacial development is severely disrupted in 5 dpf *polr1c^hi1124/hi1124^* zebrafish (Noack Watt et al., 2016), hereafter referred to as *polr1c^-/-^.* The interdependence of craniofacial and neural development during embryogenesis has long been appreciated, and the co-occurrence of facial and brain anomalies in congenital disorders gave rise to the concept that “the face predicts the brain” (DeMyer et al., 1964). To determine if neuronal development was also affected in *polr1c^-/-^* zebrafish, we examined neuronal populations using immunofluorescence (IF) staining against the pan-neuronal marker HuC (Kim et al., 1996) (Fig. 1 A,B). HuC-positive cells were detected throughout the head, including in the cranial ganglia and the enteric nervous system (ENS). Quantification of HuC+ volume throughout the head at 5 dpf revealed no statistically significant differences between control and *polr1c^-/-^* larvae (Fig. 1 C, region defined in A’, B’); however, the distribution of HuC staining in the forebrain and midbrain appeared altered in mutants comparted to controls.

**Fig 1.**
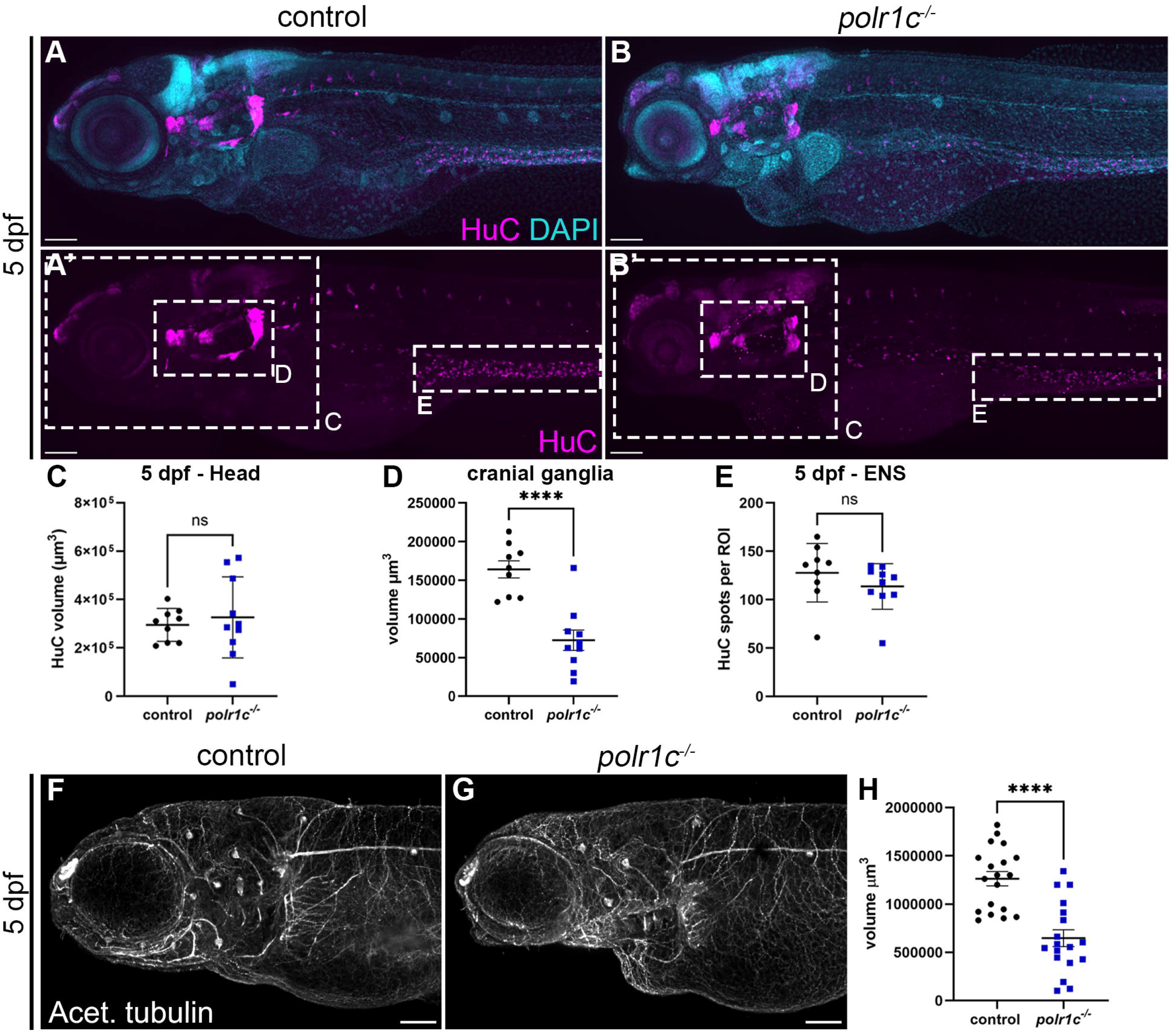
Reduced neuronal development and axons are observed in *polr1c^-/-^* zebrafish at 5 dpf. A,B) Staining against pan-neuronal marker HuC (magenta) and nuclei (DAPI, cyan) show differences in head morphology and HuC distribution. *polr1c^-/-^*larvae show an overall reduction in the size of the head, with a more striking reduction in the midbrain region. **C)** Quantification of HuC throughout the head did not reveal significant differences between control and mutant larvae. **D)** Quantification of the region of the cranial ganglia revealed a significant reduction in this area ****p<0.0001. **E)** Quantification of HuC in the enteric nervous system (ENS) did not reveal significant differences. n = 9 controls, 10 mutants. **F,G)** Immunofluorescence staining for acetylated tubulin shows differences in axon branching in *polr1c^-/-^* larvae, with mutants showing fewer large axons (arrows). **H)** Quantification of acetylated tubulin staining shows a significant reduction in overall volume throughout the head. This is likely a reflection of fewer large diameter axons. ****p<0.0001 n = 19 controls, 19 mutants. Graphs represent the mean ± s.d., and p values were determined using a two-tailed Student’s t-test.

We previously demonstrated that the NCC population is reduced in *polr1c^-/-^*embryos (Noack Watt et al., 2016), which contributes to the cranial ganglia and ENS, so we next examined these populations individually. The volume of the cranial ganglia was significantly reduced in *polr1c^-/-^* larvae (Fig. 1D, region defined in A’, B’), and both placode-derived ganglia and ganglia of mixed NCC and placode origins are affected. This indicates that neuronal differences may extend beyond NCC-derived tissues. In contrast, we did not detect a difference in the extent of HuC staining nor the number of HuC positive cells within the ENS (Fig. 1E, region defined by A’, B’). However, it is possible that there are differences in the ENS that we did not detect with our limited examination.

We next assessed axonal projections via IF staining for acetylated tubulin (Wilson et al., 1990). Qualitatively, the axonal projections appeared less organized in *polr1c^-/-^*zebrafish, with increased branching and fewer large diameter axons compared to controls (Fig. 1 F,G). Consistent with these observations, the total volume of acetylated tubulin was significantly reduced in *polr1c* mutants (Fig. 1H). Altogether, these results demonstrate that loss of *polr1c* disrupts the development of the cranial ganglia and axons, with both NCC-derived and non-NCC derived structures affected.

### Loss of *polr1c* disrupts myelin development and reduces oligodendrocyte migration

Given the altered axonal organization in *polr1c^-/-^* larvae (Fig. 1) and the association of pathogenic variants in *POLR1C* with hypomyelinating leukodystrophy (Gauquelin et al., 2019; Thiffault et al., 2015), we next examined myelin development. In zebrafish, oligodendrocytes begin terminal differentiation and initiate expression of myelin-associated genes, including myelin basic protein (*mbp)*, at about 72 hours post fertilization (hpf). We used *Tg(mbp:tagRFP),* marking cells positive for *myelin basic protein a* (Hines et al., 2015), which is a major component of the myelin sheath (de Monasterio-Schrader et al., 2012), and *Tg(olig2:egfp),* marking cells positive for *oligodendrocyte transcription factor 2* (Shin et al., 2003), to visualize myelinating cells in the CNS (Fig. 2 A,B).

**Fig 2.**
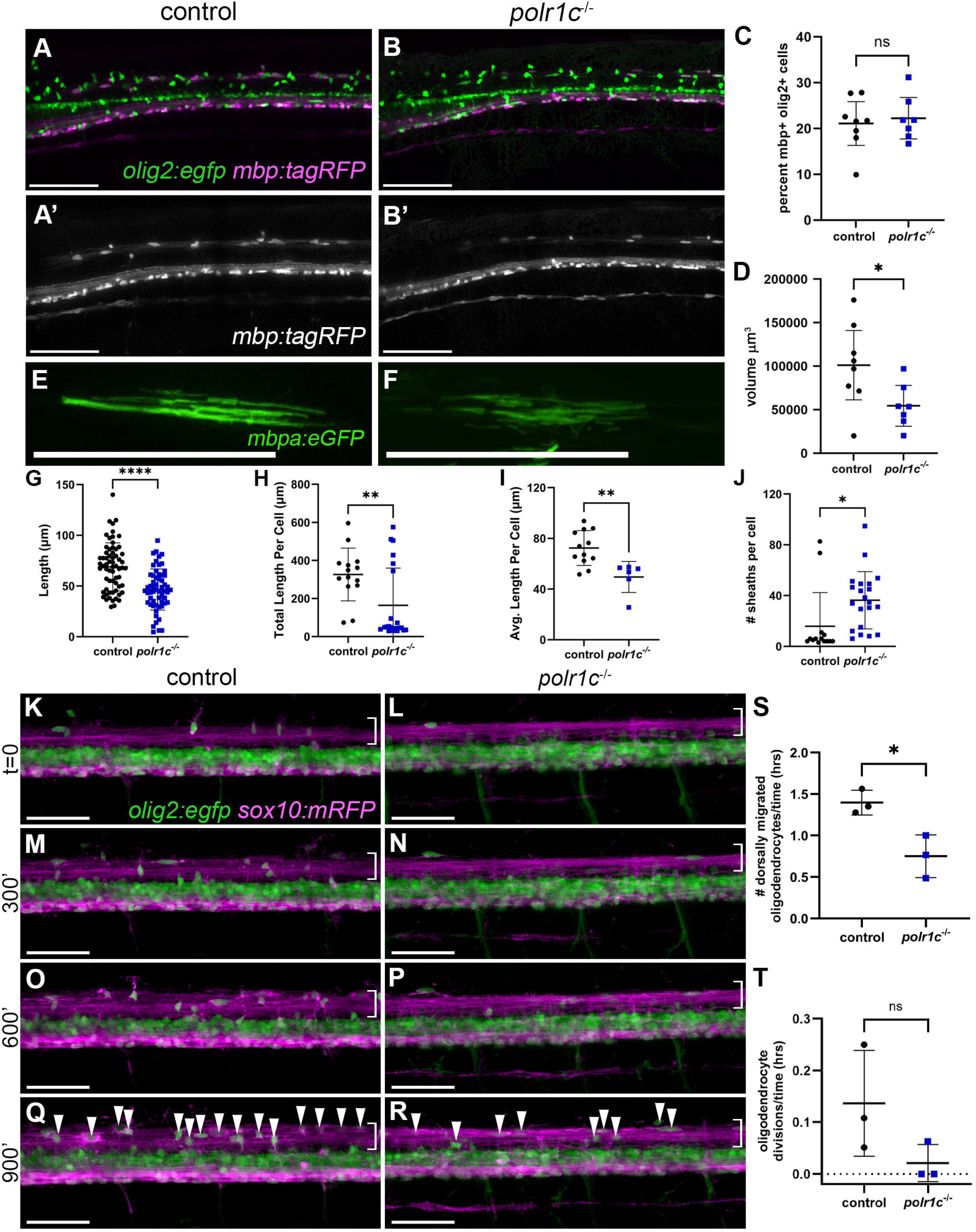
Myelination is reduced in the spinal cord of *polr1c^-/-^* embryos at 5 dpf. A-D) Expression of *olig2:egfp* and *mbp:tagRFP* in control and *polr1c^-/-^*embryos reveals no change in the percentage of *olig2+mbp+* cells between control and mutants (C); however, a significant reduction in overall *mbp:tagRFP* volume was detected in mutant zebrafish (D), indicative of reduced myelination. *p = 0.045. n = 8 controls, 7 mutants. **E-J)** An *mbp:egfp-CAAX* plasmid was injected into control and *polr1c^-/-^* embryos at the 1-cell stage to induce mosaic expression and myelin sheaths were quantified at 5 dpf in the dorsal spinal cord. Myelin sheaths appeared shorter overall in *polr1c^-/-^* zebrafish relative to controls (E, F) and quantification revealed reductions in the overall sheath length (G; n = 64 control sheaths, 59 *polr1c*^-/-^ sheaths ****p<0.0001), total sheath length per cell (H; n = 12 controls, 6 *polr1c^-/-^;* **p = 0.0072), and average sheath length per cell (I; n = 12 controls, 6 *polr1c^-/-^;* **p = 0.0042). The total number of sheaths per cell was slightly increased (J; n = 12 controls, 6 *polr1c^-/-^;* *p = 0.026), indicative of more, shorter myelin sheaths per cell in *polr1c^-/-^* zebrafish. **K-T)** To determine if differences in myelin in the dorsal spinal cord were due to differences in oligodendrocyte migration, timelapse microscopy of embryos expressing *olig2:egfp* and *sox10:mRFP* was performed from 54 hpf – 72 hpf (K-R). Fewer oligodendrocytes migrated dorsally in *polr1c^-/-^* embryos (Q vs R), and quantification showed that there was less migration per hour in mutant embryos (S; *p = 0.028). To determine if this was due to a difference in proliferation, we quantified oligodendrocyte divisions and found no significant difference between controls and *polr1c^-/-^* zebrafish (T) n = 3 controls, 3 *polr1c^-/-^*. Scale bar = 100 µm. Graphs represent the mean ± s.d., and p values were determined using a two-tailed Student’s t-test.

We quantified reporter expression in the anterior spinal cord (somites 1-7) at 5 dpf to determine if there were significant differences (Fig. 2 C, D). The proportion of *olig2:egfp*+ cells that were also *mbp:tagRFP*+ did not differ between control and *polr1c^-/-^*zebrafish (Fig. 2C), indicating that the expression of *mbp* by oligodendrocytes was not impaired at this stage. However, total *mbp:tagRFP* volume was significantly reduced in *polr1c^-/-^* zebrafish, particularly in the more dorsal region of the spinal cord (Fig. 2D). This indicates that although a similar proportion of oligodendrocytes express *mbp,* the extent of myelination may be affected.

To determine if reduced *mbp:tagRFP* expression persists, we examined zebrafish at 7 dpf (Fig. S1). At this stage, both the proportion of myelinating oligodendrocytes (*mbp+ olig2+* cells, Fig. S1C) as well as the overall *mbp:tagRFP+* volume (Fig. S1D) were significantly reduced. This indicates that the myelin phenotype persists and becomes more pronounced at later developmental stages. We next examined myelin structure using transmission electron microscopy (TEM) of transverse sections of the anterior spinal cord (Fig. S1 E-H). Myelin surrounding CNS axons was moderately affected in *polr1c^-/-^* zebrafish (Fig. S1 E,F). In the peripheral nervous system (PNS), the myelin surrounding the axons was drastically reduced in *polr1c^-/-^* zebrafish (Fig. S1 G,H), suggesting that the ENS may be affected. Thus, the loss of *polr1c* in zebrafish may disrupt myelination in both the CNS and PNS.

To better quantify individual myelin sheaths within the developing CNS, we injected a *mbpa:egfp-CAAX* construct at the one-cell stage and then analyzed individual *egfp+* cells at 5 dpf in the dorsal region of the spinal cord (Fig 2. E-J). Myelinating cells in *polr1c^-/-^* zebrafish formed significantly shorter myelin sheaths compared to controls (Fig. 2G). The sum of myelin sheath length per cell (Fig. 2H) and the average overall sheath length per cell (Fig. 2I) were reduced in *polr1c^-/-^* zebrafish. In contrast, there was an increased number of sheaths per cell (Fig. 2J), suggesting that mutants form an increased number of short myelin sheaths. These findings indicate that *polr1c* alters myelin sheath organization.

Reduced *mbp:tagRFP* expression was consistently most apparent in the dorsal spinal cord, so we next examined whether oligodendrocyte migration to this region was affected in *polr1c^-/-^* zebrafish. We used timelapse imaging of zebrafish expressing *olig2:egfp* and *sox10:mRFP* (Kucenas et al., 2008), to label oligodendrocytes and track their migration from 2 to 3 dpf (Fig. 2 K-T). At t=0 (54 hpf), a small number of dorsally located oligodendrocytes (*olig2+; sox10+*) were present in both control and *polr1c^-/-^* zebrafish (Fig. 2 K, L; bracket). During the next 15 hours, we observed dorsal migration of individual oligodendrocytes (Fig. 2 M-R). However, fewer cells underwent this dorsal migration in *polr1c^-/-^* embryos, resulting in fewer oligodendrocytes in the dorsal spinal cord (arrowheads Fig. 2 Q,R; Fig. 2S). Some of the dorsally migrated cells underwent cell division during imaging; however, the number of divisions over time did not significantly differ between controls and *polr1c^-/-^* embryos (Fig. 2T). This suggests that while there is a deficiency in the number of dorsally migrated oligodendrocytes, it is not due to differences in their proliferation at this stage. Importantly, the oligodendrocytes that do migrate in *polr1c^-/-^* embryos reached the appropriate dorsal location, demonstrating that the capacity to migrate is preserved in *polr1c* mutants. Altogether, these data indicate that loss of *polr1c* disrupts myelin sheath growth and reduces the number of oligodendrocytes that migrate to the dorsal spinal cord.

### *olig2* expression is altered in *polr1c^-/-^* embryos

Timelapse imaging revealed reduced *olig2:egfp* expression in *polr1c^-/-^* embryos, suggesting that differences in the *olig2+* population might precede the reduction in oligodendrocyte migration (Fig. 2). Olig2 is expressed in the pMN domain of the developing spinal cord, which gives rise to motor neurons and oligodendrocytes (Park et al., 2002). In the head, *olig2* is expressed in the neurosecretory preoptic area, posterior tuberculum, dorsal thalamus, and hindbrain (Shin et al., 2003; Zannino and Appel, 2009). To define when differences in *olig2* expression emerge, we examined *olig2:egfp* from 26 hpf through 74 hpf (Fig. 3 A-H). This time period covers the expression of *olig2* in motor neuron progenitors (prior to 30 hpf), in oligodendrocyte precursor cells (after 30 hpf), in pre-myelinating oligodendrocytes (48 hpf), and in myelinating oligodendrocytes (72 hpf) (Ackerman and Monk, 2016).

**Fig 3.**
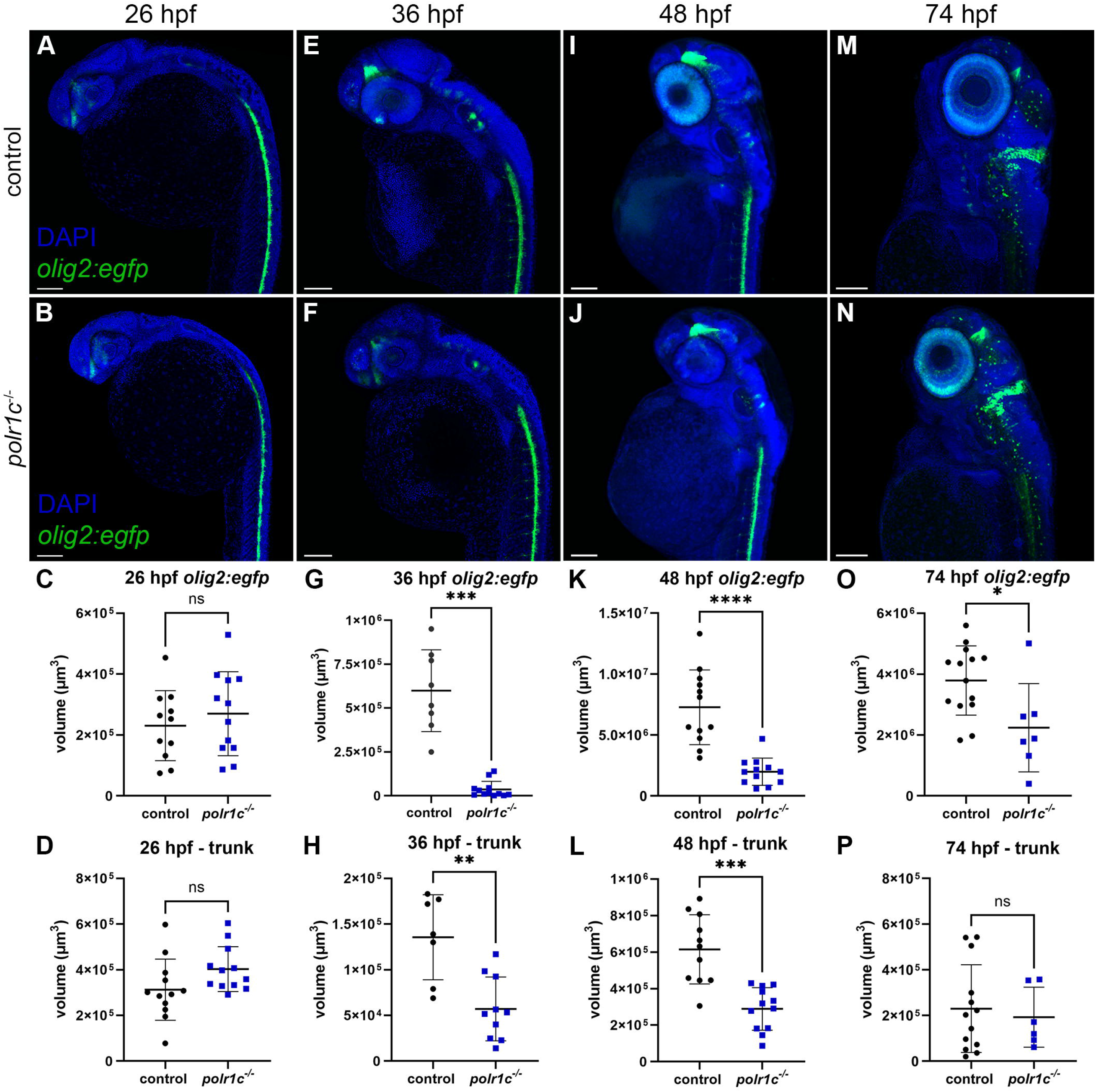
*olig2:gfp* expression is reduced in *polr1c^-/-^* embryos from 36 hpf. Control and *polr1c^-/-^*zebrafish were examined from 26 hpf - 74 hpf to determine the onset of differences in *olig2:egfp* expression. **A, B)** At 26 hpf, no significant differences were detected in *olig2:egfp* expression in the head (C) or trunk (D) of *polr1c^-/-^* embryos. n = 12 controls, 12 mutants. ns, not significant. **E,F)** By 36 hpf, significant reductions were detected in both the head (G) and trunk (H). n = 7 controls, 10 mutants. ***p = 0.0002 **p = 0.0033. **I,J)** At 48 hpf, these reductions persist in both the head (K) and trunk (L). ****p < 0.0001; ***p = 0.0001. n = 12 controls, 12 mutants. **M,N)** At 72 hpf, olig2 expression remains reduced in the head (O), which is notably smaller in *polr1c^-/-^* zebrafish at this stage. *p = 0.033. However, expression in the trunk (P) was not significantly different. n = 14 controls, 7 mutants. Scale bar = 100 µm. Graphs represent the mean ± s.d., and p values were determined using a two-tailed Student’s t-test.

At 26 hpf, *olig2:egfp* expression in both the head and the trunk was similar between control and *polr1c^-/-^* embryos, with no significant differences detected (Fig. 3 A-D). This suggests that the expression of *olig2* in the pMN domain is maintained during motor neuron development. By 36 hpf, a significant reduction in *olig2:egfp* expression was evident in both the head and the trunk (Fig. 3 E-H). In the head, reductions in *olig2:egfp* are most apparent in the forebrain and hindbrain. At 48 hpf, reduced *olig2:egfp* expression persists in both the head and trunk in the *polr1c^-/-^* zebrafish (Fig. 3 I-L) and remains reduced in the head at 74 hpf (Fig. 3 M-O). In contrast, there was no longer a statistically significant difference in *olig2:egfp* expression observed in the trunk at 74 hpf (Fig. 3P). These data identify reduced *olig2:egfp* expression occurs between 26 and 36 hpf, preceding the reduction in oligodendrocyte migration.

The reduction in *olig2:egfp* expression at 36 hpf, which coincides with the neurogenesis to gliogenesis switch, led us to hypothesize that there may be differences in neuronal populations, including those derived from the pMN, at 36 hpf. Quantification of pan-neuronal maker HuC (Kim et al., 1996) at 36 hpf revealed no significant differences in overall volume between control and *polr1c^-/-^* embryos (Fig. S2 A, B), consistent with the absence of major differences in HuC volume at 5 dpf (Fig. 1). We next examined Isl1, which is a marker of motor neurons that are derived from the pMN domain, as well as Rohon-Beard cells and the cranial ganglia (Inoue et al., 1994; Korzh et al., 1993). Isl1 expression in the head was not significantly different between control and *polr1c^-/-^* embryos (Fig. S2 C-E); however, Isl1 in the trunk was significantly reduced in *polr1c^-/-^* embryos (Fig. S2F). This indicates that developmental differences in the spinal cord are evident by 36 hpf, affecting both the pMN-derived motor neuron population and the emerging oligodendrocyte lineage. The reduction in *olig2:egfp* expression before oligodendrocyte migration suggests that this earlier reduction could influence the number of oligodendrocytes that subsequently migrate to the dorsal spinal cord.

### Cell death is increased in the dorsal neural tube of *polr1c^-/-^* embryos

We next hypothesized that the reduction in *olig2:egfp* expression in *polr1c^-/-^* embryos at 36 hpf was due to increased apoptosis at an earlier stage. Previous studies revealed increased cell death in *polr1c^-/-^*zebrafish (Noack Watt et al., 2016), and impaired rRNA transcription can induce apoptosis in neural progenitors (Parlato et al., 2008) and post-mitotic neurons (Kalita et al., 2008). We assessed cell death using TUNEL staining and proliferation using IF against phospho-histone H3 (pHH3) at 26 hpf.

Increased TUNEL staining was observed throughout the head and neuroepithelium of *polr1c^-/-^* embryos compared to controls (Fig. 4 A-E); however, pHH3 was unchanged at this stage (Fig. 4F). Analysis of transverse sections through the neural tube revealed that cell death is localized primarily to the dorsal region of the neural tube (Fig. 4D’), where NCC progenitors reside (Knecht and Bronner-Fraser, 2002). Surprisingly, we did not observe cell death in the pMN domain or nearby domains, where OPCs are located (Ravanelli and Appel, 2015). These findings suggest that the subsequent reduction in *olig2:egfp* expression and impaired myelin development in *polr1c* mutants cannot be explained simply by the loss of pMN-derived progenitors.

**Fig 4.**
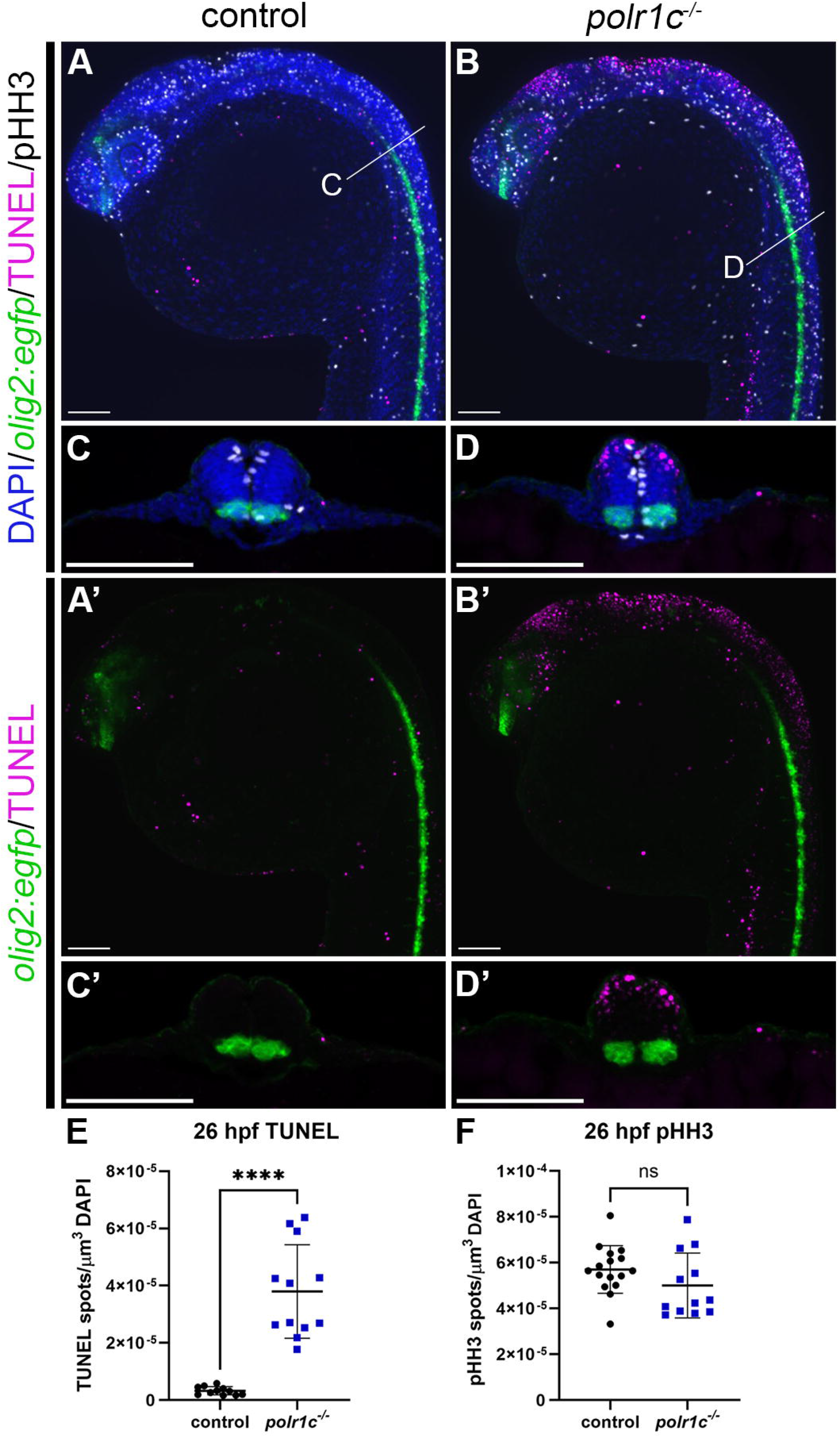
Cell death is increased in the dorsal neural tube at 26 hpf. A, **B)** Control and *polr1c^-/-^*embryos expressing *Tg(olig2:egfp)* were stained for TUNEL (magenta) as a marker of cell death, pHH3 (white) as a marker of proliferation, DAPI (blue) for all nuclei. Increased cell death was observed in the head and neural tube. **C, D)** Cross sections through the neural tube reveal increased cell death in the dorsal portion of the neural tube, but not in the *olig2:egfp* domain. pHH3 positive cells are detected in the *olig2:egfp* domain in both control and mutant zebrafish. **A’-D’)** Images of TUNEL and *olig2:egfp* only confirm the absence of TUNEL+ cells in the *olig2* domain. **E, F)** Quantification reveals increased cell death in *polr1c^-/-^* embryos at 26 hpf (E; ****p < 0.0001), while proliferation remains unchanged (F; ns, not significant). n = 16 controls, 12 polr1c^-/-^. Scale bar = 100 µm. Graphs represent the mean ± s.d., and p values were determined using a two-tailed Student’s t-test.

### The Tp53 pathway is upregulated in OPCs and NCCs in *polr1c^-/-^* embryos

To determine the transcriptional changes that preceded reduced *olig2:egfp* expression, occurring at the onset of the mutant phenotype, we performed bulk RNA sequencing of control and *polr1c^-/-^* embryos at 24 hpf. Differential expression analyses revealed there are already widespread transcriptional changes in *polr1c* mutants at this stage, with a greater number of upregulated genes compared to those showing downregulation (Fig. 5A). Among the upregulated genes were multiple targets in the Tp53 pathway, including *tp53; mdm2*, a negative regulator of Tp53 (Haupt et al., 1997; Honda et al., 1997; Kubbutat et al., 1997); and *cdkn1a*, a downstream target of Tp53 (Löhr et al., 2003) (Fig. 5A). Gene Ontology (GO) analysis of significantly upregulated genes revealed enrichment of terms related to Tp53 signaling, cell cycle regulation, and DNA damage (Fig. 5B), all of which have been previously implicated in TCS (Jones et al., 2008; Noack Watt et al., 2016; Sakai et al., 2016).

**Fig 5.**
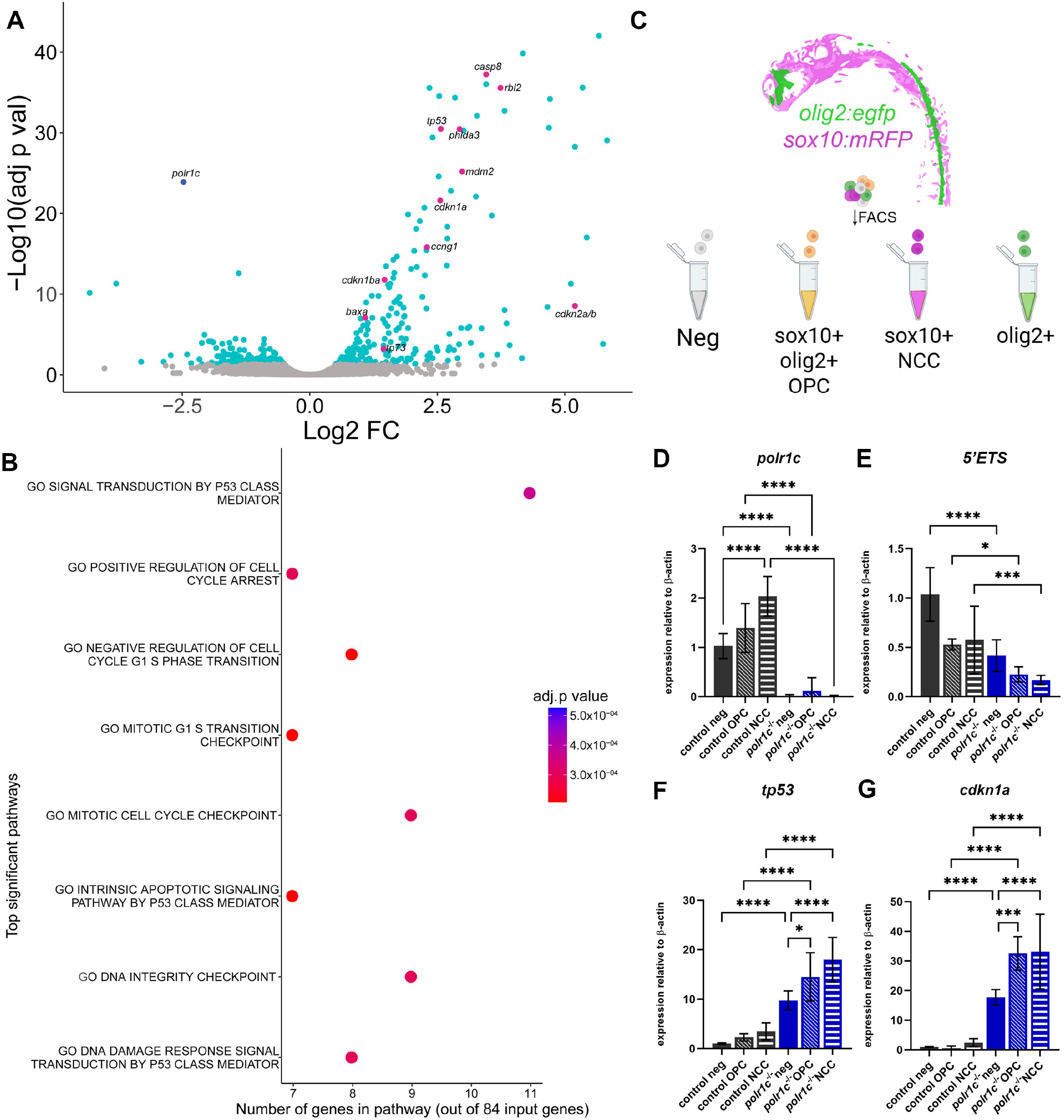
RNA-sequencing reveals upregulation of the Tp53 pathway at 24 hpf in *polr1c^-/-^* zebrafish. **A)** Volcano plot of differences in gene expression between control (*polr1c^+/+^*) and *polr1c^-/-^*embryos at 24 hpf. *polr1c* was significantly downregulated (dark blue), while several genes involved in cell cycle regulation and cell death were significantly upregulated (magenta). **B)** GO term analysis of the significantly upregulated genes in *polr1c^-/-^*embryos revealed increased Tp53-dependent signaling and cell cycle arrest. **C)** Schematic of cell sorting scheme. Embryos expressing *Tg(olig2:egfp)* and *Tg(sox10:mRFP)* were dissociated and sorted at 32 hpf. *sox10+olig2+* cells represent the oligodendrocyte progenitors (OPCs) while *sox10+* only cells represent neural crest cells (NCC). RNA from the sorted cells was extracted for qRT-PCR experiments in D-G. **D)** qRT-PCR for *polr1c* confirms downregulation of *polr1c* in mutant embryos across all cell types (****p < 0.0001). In controls, *polr1c* is more highly expressed in NCC relative to negative cells. **E)** 5’ETS pre-rRNA is downregulated in all cell types in *polr1c^-/-^*embryos. ****p<0.0001; *p = 0.021; ***p = 0.0007. **F)** *tp53* is significantly increased in all cell types in *polr1c^-/-^* embryos. The OPCs and NCCs express a relatively higher level of *tp53* compared to negative cells in *polr1c^-/-^* embryos. ****p<0.0001; *p = 0.0143. **G)** Consistent with high *tp53* expression, *cdkn1a* expression is higher in all *polr1c^-/-^* cells relative to controls, with the highest levels detected in OPCs and NCCs. ****p<0.0001; ***p = 0.0004. Graphs represent the mean ± s.d., and p values were determined using a one-way ANOVA with Tukey’s multiple comparisons test.

To determine if Tp53-dependent responses occur broadly or are specific to different embryonic cell populations such as NCCs and OPCs, we used fluorescence activated cell sorting (FACS) to isolate these populations from embryos expressing *olig2:egfp* and *sox10:mRFP* at 32 hpf (Fig. 5C). *olig2:egfp* and *sox10:mRFP* double positive cells were isolated to represent OPCs, while *sox10:mRFP* positive and *olig2:egfp* negative cells represent the NCC population, and cells negative for both reporters represent a broader embryonic population. Quantitative RT-PCR (qRT-PCR) confirmed reduced *polr1c* expression in each mutant cell population (Fig. 5D). Interestingly, in controls, we observed that *polr1c* expression was higher in NCCs (control NCC) compared to the reporter negative population (control neg), consistent with our previous *in situ* data suggesting that *polr1c* is enriched in NCCs (Noack Watt et al., 2016). To determine if the reduction in *polr1c* also impaired rRNA transcription by Pol I, we examined expression of the 5’ETS (external transcribed spacer) of pre-rRNA as a proxy for nascent rRNA transcription. 5’ETS expression was reduced in all *polr1c* mutant cell populations examined, including OPCs and NCCs (Fig. 5E), demonstrating that rRNA transcription is impaired across multiple cell types.

Disruptions in rRNA transcription and ribosome biogenesis are known to trigger Tp53-dependent cell death (Donati et al., 2013; Rubbi and Milner, 2003; Sloan et al., 2013). To assess Tp53 activation across cell types, we examined the expression of *tp53* and its downstream target *cdkn1a*.

Both genes were significantly upregulated in OPCs, NCCs, and reporter negative cells in *polr1c^-/-^* embryos compared to controls (Fig. 5 F,G). Intriguingly, NCC and OPCs both exhibited higher levels of *tp53* and *cdkn1a* compared to the reporter negative population. However, this enhanced Tp53 response has different cellular outcomes in the two populations (Fig. 4). The NCC progenitors in the dorsal neuroepithelium exhibit extensive apoptosis, which we did not detect in the pMN domain despite evidence of significant Tp53 pathway activation. Overall, these data demonstrate that *polr1c* loss-of-function reduces rRNA transcription across embryonic cell types and results in broad activation of Tp53 signaling. Tp53 pathway activation is particularly strong in NCCs and OPCs, but cell death occurs in a more tissue-specific manner.

### Tp53 inhibition in *polr1c^-/-^* zebrafish reduces cell death and leads to slight improvements in *olig2:egfp* expression

Given the broad activation of Tp53 signaling in *polr1c* mutants, including significant upregulation of *tp53* and *cdkn1a* in OPCs (Fig. 5), we hypothesized that inhibition of *tp53* could reduce cell death and ameliorate developmental anomalies in mutant embryos, including in oligodendrocytes. To test this, we genetically inhibited *tp53* and assessed cell death (TUNEL), proliferation (pHH3), and *olig2:egfp* expression at 26 hpf, 36 hpf, and 72 hpf (Fig. 6).

**Fig 6.**
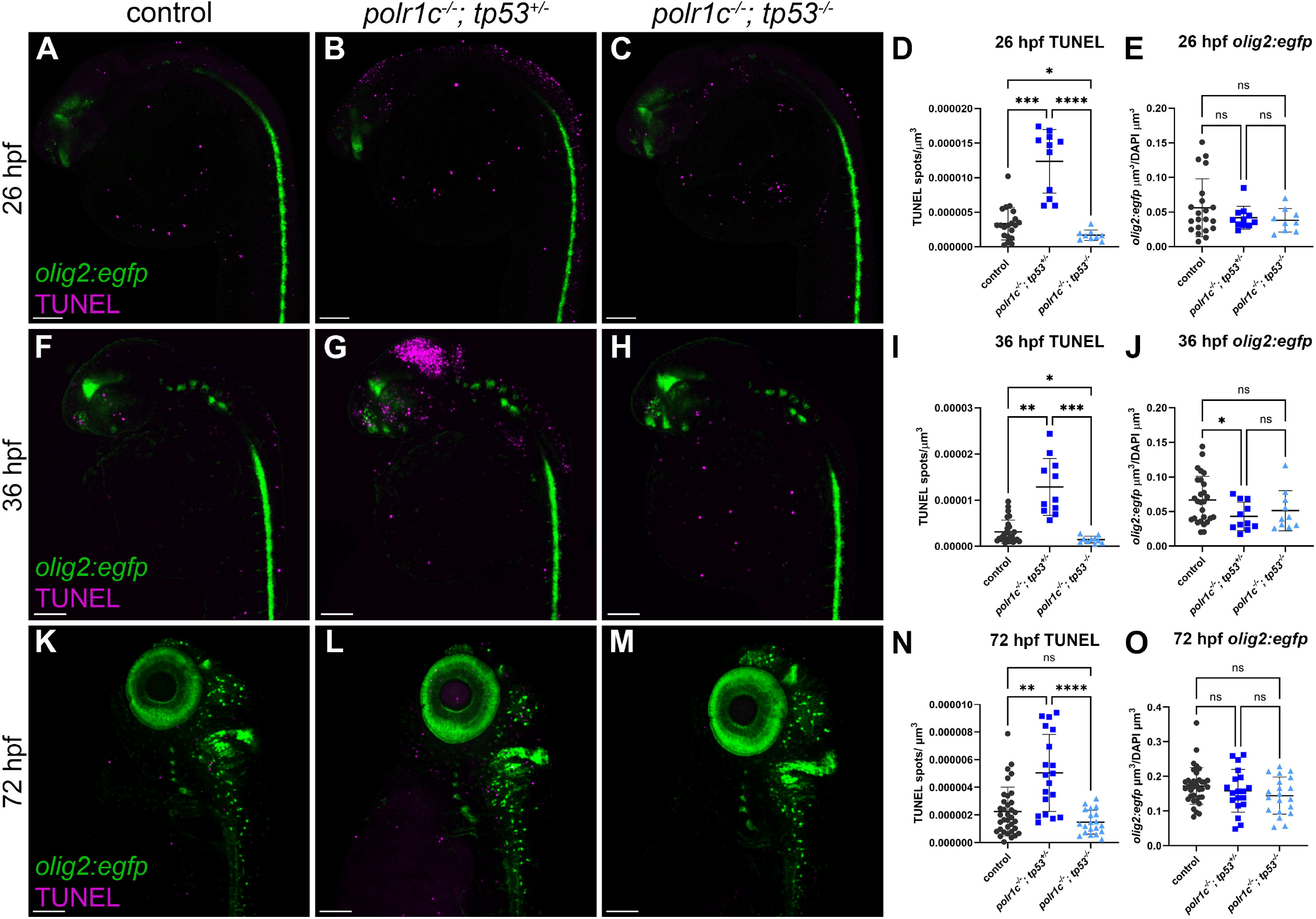
*tp53* inhibition reduces cell death and slightly improves *olig2:egfp* expression. Cell death was assessed at 26, 36, and 72 hpf in control and *polr1c^-/-^* zebrafish expressing *Tg(olig2:egfp).* **A-E)** At 26 hpf, cell death is significantly reduced in *polr1c^-/-^; tp53^-/-^*embryos (C) compared to *polr1c^-/-^; tp53^+/-^* embryos (B) and controls (A). Quantification in D. ***p = 0.0001; *p = 0.02; ****p<0.0001. *olig2:egfp* expression is unchanged at this stage, with no significant differences across groups (E). n = 20 controls, 11 *polr1c^-/-^; tp53^+/-^,* 9 *polr1c^-/-^; tp53^-/-^.* **F-J)** At 36 hpf, cell death is significantly reduced in *polr1c^-/-^; tp53^-/-^* embryos (H) compared to *polr1c^-/-^; tp53^+/-^*embryos (G) and *tp53^+/-^* controls (F). Quantification in I. **p = 0.001; *p = 0.01; ***p = 0.0003. *olig2:egfp* expression is reduced in *polr1c^-/-^; tp53^+/-^* embryos relative to controls (*p = 0.040), while in *polr1c^-/-^; tp53^-/-^* embryos, expression shows some improvement and is not significantly different from either controls or *polr1c^-/-^; tp53^+/-^* embryos. n = 27 controls, 11 *polr1c^-/-^; tp53^+/-^,* 10 *polr1c^-/-^; tp53^-/-^.* **K-O)** At 3 dpf, cell death is significantly reduced in *polr1c^-/-^; tp53^-/-^* embryos (M) compared to *polr1c^-/-^; tp53^+/-^*embryos (L) and not significantly different from controls (K). Quantification in N. ***p = 0.0007; ****p<0.0001. *olig2:egfp* expression is unchanged at this stage, with no significant differences across groups (O). n = 34 controls, 19 *polr1c^-/-^; tp53^+/-^,* 20 *polr1c^-/-^; tp53^-/-^.* Scale bar = 100 µm. Graphs represent the mean ± s.d., and p values were determined using a one-way ANOVA with Tukey’s multiple comparisons test.

At 26 hpf, *polr1c^-/-^; tp53^-/-^* embryos exhibited a significant reduction in cell death compared to *polr1c^-/-^; tp53^+/-^* embryos (Fig. 6 A-D; Fig. S3), confirming that much of the apoptosis associated with the loss of *polr1c* is Tp53-dependent. Consistent with our previous observations, *olig2:egfp* expression was not significantly different at this stage (Fig. 6E). At 36 hpf, increased cell death was evident in the brain of *polr1c^-/-^; tp53^+/-^* embryos, but was nearly absent in *polr1c^-/-^; tp53^-/-^* embryos (Fig. 6 F-I; Fig. S4). Interestingly, *polr1c^-/-^; tp53^+/-^* embryos showed an intermediate *olig2:egfp* expression phenotype at this stage. While *polr1c^-/-^; tp53^+/-^* embryos exhibited a significant reduction in *olig2:egfp* expression compared to controls, *olig2:egfp* expression in *polr1c^-/-^; tp53^-/-^* embryos was not significantly different from either controls or *polr1c^-/-^; tp53^+/-^* embryos (Fig. 6J). The overall morphology of *polr1c^-/-^; tp53^-/-^* embryos was improved relative to *polr1c^-/-^; tp53^+/-^* embryos (Fig. S4), consistent with the broad benefit of *tp53* inhibition, but suppression of Tp53-dependent cell death resulted in only a modest improvement in the *olig2:egfp* population. By 72 hpf, apoptosis remained significantly reduced in *polr1c^-/-^; tp53^-/-^* zebrafish compared to *polr1c^-/-^; tp53^+/-^*zebrafish and was not significantly different from controls (Fig. 6 K-N; Fig S5). *olig2:egfp* expression was also not significantly different across groups at this stage (Fig. 6O).

To determine if differences in proliferation could partially account for the observed differences across stages, we quantified pHH3 throughout the anterior region of the embryos at each developmental stage. No significant differences were detected across groups at 26, 36, or 72 hpf (Fig. S6). Although this analysis does not exclude the possibility that there are cell type-specific changes in proliferation in the *olig2+* population, it indicates that loss of *tp53* does not produce a detectable global change in proliferation at the stages examined (Figs. S3-S6). Altogether, these data demonstrate that Tp53-dependent apoptosis in *polr1c^-/-^* embryos occurs at multiple stages of development (Fig. 6 D,I,O), and that inhibition of *tp53* reduces cell death and improves overall embryonic morphology. In contrast, the effect on *olig2:egfp* expression is modest, suggesting that differences in the *olig2+* population cannot be explained by Tp53-dependent cell death and there may be other Tp53-independent factors involved.

### *tp53* inhibition does not improve myelin development in *polr1c* mutant zebrafish

Genetic inhibition of *tp53* reduced apoptosis and improved overall development in *polr1c* mutants, so we next asked if these improvements extended to myelin development. We imaged the anterior spinal cord of control, *polr1c^-/-^; tp53^+/+^*, *polr1c^-/-^; tp53^+/-^*, and *polr1c^-/-^; tp53^-/-^* zebrafish expressing *mbp:tagRFP* and *olig2:egfp* at 5 dpf (Fig. 7 A-D). The relative volume of *olig2:egfp* was increased in *polr1c^-/-^*larvae following *tp53* inhibition (Fig. 7E), consistent with the modest improvement in *olig2:egfp* we observed at earlier stages (Fig. 6). However, *mbp:tagRFP* volume remained reduced in *polr1c^-/-^; tp53^-/-^* zebrafish compared to controls and was not significantly different from *polr1c^-/-^; tp53^+/+^*siblings (Fig. 7F). Consequently, the proportion of *olig2:egfp*+ cells that were also *mbp:tagRFP*+ in *polr1c^-/-^; tp53^-/-^* zebrafish was reduced compared to controls (Fig. 7G). These results demonstrate that although *tp53* inhibition improves the *olig2*+ population, it does not improve *mbp* expression. This indicates that impaired myelin development in *polr1c* mutants may occur through persistent molecular consequences of *polr1c* loss, encompassing *tp53-*independent mechanisms.

**Fig 7.**
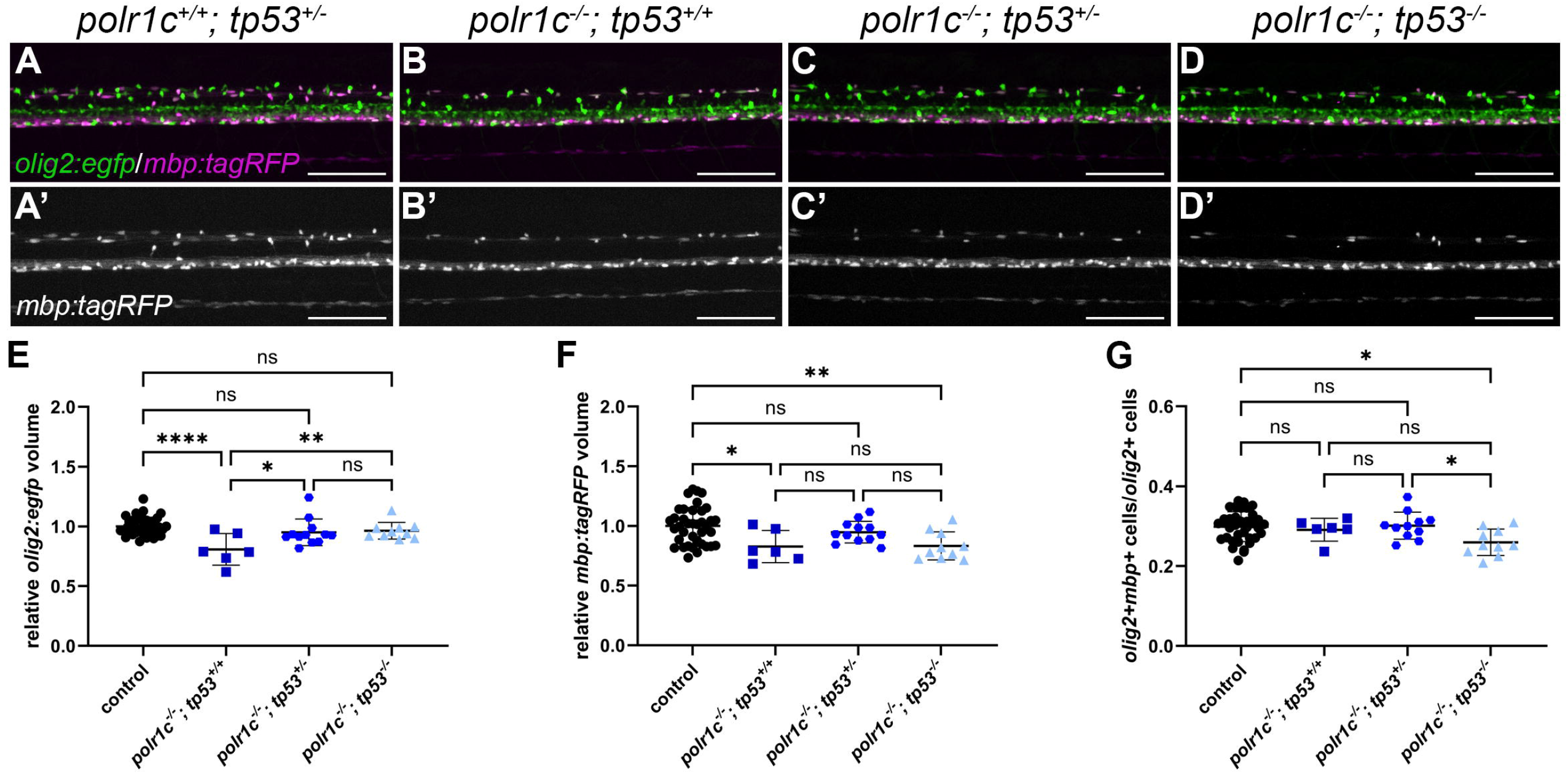
*mbp:tagRFP* expression remains reduced in *polr1c^-/-^; tp53^-/-^* zebrafish at 5 dpf. **A-D)** *olig2:egfp* (green) and *mbp:tagRFP* (magenta) expression in the spinal cord of 5 dpf zebrafish. *mbp:tagRFP* expression (A’-D’) remains reduced in *polr1c^-/-^* zebrafish regardless of *tp53* status. **E)** Quantification of the relative *olig2:egfp* volume shows a significant improvement in *olig2:egfp* in *polr1c^-/-^; tp53^+/-^* and *polr1c^-/-^; tp53^-/-^* zebrafish. ****p<0.0001; *p = 0.010; **p = 0.0056; ns, not significant. **F)** Quantification of the relative *mbp:tagRFP* volume. *mbp:tagRFP* expression is not improved in *polr1c^-/-^; tp53^-/-^* zebrafish compared to *polr1c^-/-^; tp53^+/+^* zebrafish and remains reduced compared to controls. *p = 0.031; **p = 0.0066; ns, not significant. **G)** Quantification of the proportion of *olig2:egfp; mbp:tagRFP* double positive cells over the total number of *olig2:egfp* positive cells. This ratio was significantly reduced in *polr1c^-/-^; tp53^-/-^* zebrafish relative to controls (*p = 0.015), consistent with partial rescue of *olig2:egfp*, but not *mbp:tagRFP* expression. *polr1c^-/-^; tp53^-/-^* zebrafish also showed significant reduction relative to *polr1c^-/-^; tp53^+/-^* (*p = 0.045). n = 37 controls, 6 *polr1c^-/-^; tp53^+/+^;* 11 *polr1c^-/-^; tp53^+/-^;* 10 *polr1c^-/-^; tp53^-/-^.* Scale bar = 100 µm. Graphs represent the mean ± s.d., and p values were determined using a one-way ANOVA with Tukey’s multiple comparisons test.

### Pol I-dependent rRNA transcription remains impaired in *polr1c* mutant zebrafish following *tp53* inhibition

As Polr1c is a shared subunit between Pol I and Pol III, persistent defects in rRNA or tRNA production could contribute to impaired myelination independent of Tp53. We therefore assessed the expression of genes transcribed by Pol I and Pol III at 3 dpf, coinciding with the onset of myelination. *polr1c* transcript levels are nearly undetectable in *polr1c* mutants, irrespective of *tp53* status (Fig. 8A). Consistent with impaired Pol I activity, 5’ETS pre-rRNA levels were reduced by approximately 50% in *polr1c^-/-^* zebrafish and were not restored by reduction or loss of *tp53* (Fig. 8A). Thus, genetic *tp53* inhibition suppresses some of the downstream consequences of ribosomal stress without restoring the underlying defect in rRNA transcription.

**Fig 8.**
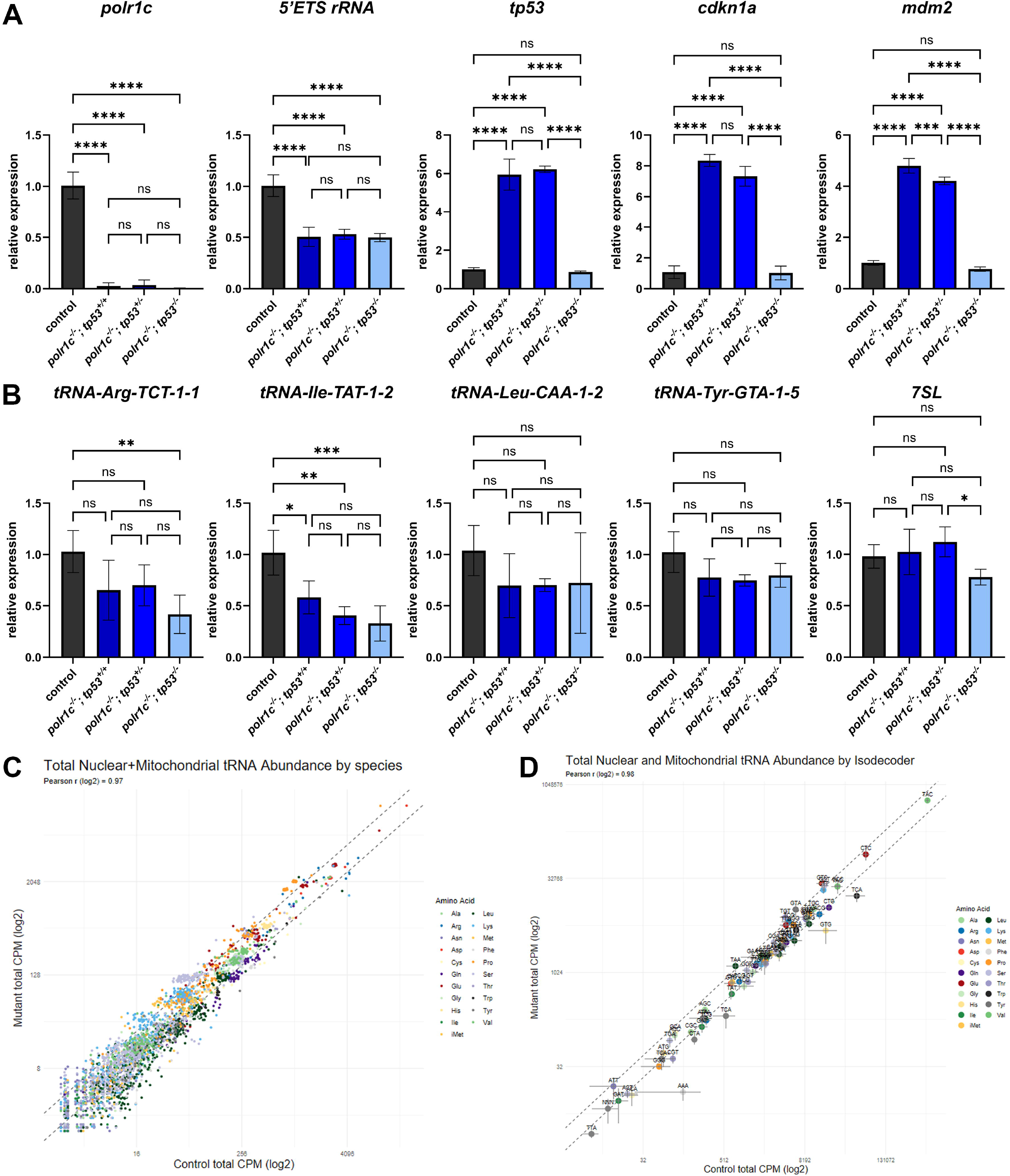
*tp53* inhibition does not improve RNA Polymerase I or III-mediated transcription at 3 dpf in *polr1c^-/-^* zebrafish. **A)** Assessment of *polr1c*, Pol I-transcribed pre-rRNA (5’ETS) and Tp53 pathway genes (*tp53, cdkn1a*, and *mdm2*) by qRT-PCR at 3 dpf. *polr1c* and *5’ETS* remain reduced in *polr1c^-/-^* zebrafish regardless of *tp53* status (****p,0.0001). *tp53, cdkn1a,* and *mdm2* are significantly upregulated in *polr1c^-/-^; tp53^+/+^*and *polr1c^-/-^; tp53^+/-^* zebrafish, while expression is reduced to wild-type levels in *polr1c^-/-^; tp53^-/-^*zebrafish. ****p<0.0001; ***p = 0.0009. n = 9 controls, 3 *polr1c^-/-^; tp53^+/+^;* 3 *polr1c^-/-^; tp53^+/-^;* 3 *polr1c^-/-^; tp53^-/-^.* Graphs represent the mean ± s.d., and p values were determined using a one-way ANOVA with Tukey’s multiple comparisons test. **B)** qRT-PCR of Pol III-transcribed tRNA genes and 7SL RNA shows that these genes are largely unaffected by *tp53* status in *polr1c^-/-^* zebrafish. *tRNA-Arg-TCT-1-1* was significantly reduced in *polr1c^-/-^; tp53^-/-^* zebrafish compared to controls (p = 0.004), but not significantly different from *polr1c^-/-^; tp53^+/+^* or *polr1c^-/-^; tp53^+/-^* zebrafish. *tRNA-Ile-TAT-1-2* was significantly reduced in *polr1c^-/-^; tp53^+/+^*, *polr1c^-/-^; tp53^+/-^*, and *polr1c^-/-^; tp53^-/-^* zebrafish (*p=0.0185; **p=0.0013; ***p=0.0005). *tRNA-Leu-CAA-1-2, tRNA-Tyr-GTA-1-5,* and *7SL* RNA were not significantly changed in *polr1c* mutants compared to controls regardless of *tp53* status. However, a slight reduction in 7SL RNA was detected in *polr1c^-/-^; tp53^-/-^* zebrafish compared their *polr1c^-/-^; tp53^+/-^* siblings (*p=0.0347). n = 9 controls, 3 *polr1c^-/-^; tp53^+/+^;* 3 *polr1c^-/-^; tp53^+/-^;* 3 *polr1c^-/-^; tp53^-/-^.* Graphs represent the mean ± s.d., and p values were determined using a one-way ANOVA with Tukey’s multiple comparisons test. **C)** Assessment of mature tRNA by tRNA-sequencing, with individual tRNAs represented and colored according to amino acid. **D)** Data from panel C, grouped by isodecoder and colored according to amino acid. Only three isodecoders (AAA, TCA, and GTG) show a reduction in expression in *polr1c^-/-^* zebrafish relative to controls, while all others remain unchanged.

To confirm the effects of *tp53* inhibition in *polr1c* mutants, we examined the expression of *tp53, cdkn1a*, and *mdm2.* All three genes were substantially upregulated in *polr1c^-/-^; tp53^+/+^*larvae, and a similar degree of induction was observed in *polr1c^-/-^; tp53^+/-^* zebrafish (Fig. 8A). Based on these data, one functional copy of *tp53* is sufficient to activate a Tp53 transcriptional response comparable to *tp53* wild-type larvae. In contrast, the relative expression of *tp53, cdkn1a*, and *mdm2* were reduced to control levels in *polr1c^-/-^; tp53^-/-^* larvae (Fig. 8A). Altogether, these data demonstrate that while *tp53* inhibition can suppress the Tp53 pathway, it does not restore rRNA transcription. Thus, impaired Pol I-dependent rRNA transcription represents one possible Tp53-independent contributor to the persistent myelin deficit in *polr1c^-/-^* zebrafish.

### Pol III-dependent transcription and tRNAs are variably affected in *polr1c* mutants

*POLR3*-related leukodystrophy has been proposed to impair Pol III-dependent production of tRNAs, which in turn affects translation and production of myelin proteins (Lata et al., 2021). We therefore examined Pol III-dependent transcripts in *polr1c^-/-^* larvae. As we previously demonstrated that 5S rRNA levels were unchanged in *polr1c^-/-^* zebrafish (Noack Watt et al., 2016), we focused on expression of pre-tRNAs as indicators of recent transcription by Pol III. At 3 dpf, the effects on pre-tRNA abundance were more variable and generally smaller than the reduction observed for pre-rRNA (Fig. 8B). *tRNA-Ile-TAT-1-2* was significantly reduced across *polr1c* mutant genotypes, with expression ranging from approximately 60% to 40% of wild-type levels. *tRNA-Arg-TCT-1-1* was significantly reduced only in *polr1c^-/-^; tp53^-/-^* zebrafish, while expression of *tRNA-Leu-CAA-1-2* and *tRNA-Tyr-GTA-1-5* in mutants did not significantly differ from controls (Fig. 8B). Levels of *7SL* RNA, another gene transcribed by Pol III, were also unchanged (Fig. 8B). This suggests that not all genes transcribed by Pol III are equally affected in *polr1c^-/-^* zebrafish, nor are they improved by *tp53* inhibition.

To assess mature tRNA abundance more broadly, we performed tRNA-sequencing of control and *polr1c^-/-^* zebrafish at 3 dpf. Analysis of individual tRNAs identified different tRNAs with either increased or decreased abundance in *polr1c^-/-^* zebrafish (Fig. 8C). When tRNAs were grouped by isodecoder (tRNAs that decode the same codon), most showed no change or modest reductions in *polr1c* mutants (Fig. 8D). These results are consistent with the variable changes we observed for pre-tRNAs by qRT-PCR and indicate that mature tRNA abundance is relatively preserved at this developmental stage. Together, these data demonstrate the loss of *polr1c* results in a significant reduction in rRNA transcription by Pol I, whereas the Pol III-dependent transcripts examined show more modest and variable differences. While contributions from altered Pol III function on myelin development cannot be excluded in this model, impaired Pol I-dependent rRNA transcription and ribosome biogenesis are prominent and persistent consequences of *polr1c* loss at the onset of myelination.

## Discussion

Here, we show that *polr1c* is required for nervous system development and myelination in zebrafish. Loss of *polr1c* alters the development of the *olig2+* population, reduces the number of oligodendrocytes that migrate dorsally, and impairs myelin sheath growth in the CNS. Although *polr1c* mutants exhibit Tp53 pathway activation and increased cell death, genetic inhibition of *tp53* does not restore myelination. Molecular analyses revealed persistent reduction in Pol I-dependent pre-rRNA transcription and reduced expression of some Pol III-transcribed tRNAs. Together, these findings indicate that impaired myelination cannot be explained solely by Tp53-dependent progenitor loss and further suggest that persistent deficits in rRNA transcription contribute to nervous system phenotypes in *polr1c* mutant zebrafish.

### *polr1c* is required for oligodendrocyte and myelin development

Myelination was impaired in both the CNS and PNS in *polr1c^-/-^*zebrafish, with a more pronounced phenotype in the PNS (Fig S1). The prominent PNS phenotype is consistent with the previously described sensitivity of NCCs to *polr1c* loss (Noack Watt et al., 2016). In the CNS of *polr1c* mutants, oligodendrocytes formed shorter myelin sheaths, but an increased number of sheaths per cell (Fig. 2). These findings are consistent with previous work showing that siRNA-mediated knockdown of *Polr1c* in cultured mouse OPCs produces immature, poorly branched oligodendrocytes (Macintosh et al., 2023). Our temporal analyses indicated that the myelin phenotype in *polr1c* mutants is likely preceded by changes in the oligodendrocyte lineage. *olig2:egfp* expression was initially similar between controls and *polr1c* mutants at 26 hpf, but was reduced by 36 hpf (Fig. 3), prior to the onset of myelination. Fewer oligodendrocytes migrated to the dorsal spinal cord in *polr1c* mutants, although the cells that did migrate reached the appropriate location (Fig. 2). This indicates that the reduction in dorsal oligodendrocytes is not due to a loss of migratory capacity and may instead arise from earlier changes in progenitor development. Reduced Isl1 expression in the trunk at 36 hpf further indicates that neuronal populations are also affected in *polr1c* mutants (Fig. S2). These findings indicate that loss of *polr1c* affects neural development prior to myelin formation, with additional defects in myelin sheath growth emerging later in development.

Whether the later defects in myelin sheath growth in *polr1c* mutants reflect cell-autonomous requirements for *polr1c* in oligodendrocytes or secondary effects of altered neuronal development remain unresolved. Myelin sheath length and thickness are influenced by axonal properties including axon diameter (Bechler et al., 2015; Murray and Blakemore, 1980), and we observed altered axonal organization in *polr1c* mutants (Fig. 1). Non-cell autonomous effects on oligodendrocyte development and migration have been demonstrated in another zebrafish mutant associated with impaired ribosome biogenesis (Simmons and Appel, 2012). Therefore, additional studies are needed to determine the cell autonomous requirements for *polr1c* in oligodendrocytes and neurons.

### Tp53 activation has distinct consequences across developing tissues

Activation of Tp53 is a common response to disrupted ribosome biogenesis and contributes to developmental defects in multiple ribosomopathies (Dutt et al., 2011; Jones et al., 2008; McGowan et al., 2008; Noack Watt et al., 2016; Watt et al., 2018). Consistent with these studies, *polr1c* mutants exhibited increased Tp53 activation and elevated neuroepithelial cell death (Figs. 4, 5). However, the consequences of Tp53 activation differed across progenitor cell populations. Both OPCs and NCCs exhibited reduced rRNA transcription and robust induction of *tp53* and *cdkn1a* (Fig. 5), while apoptosis was largely concentrated in the dorsal neuroepithelium and was not detected in the pMN domain (Fig. 4). This suggests that distinct progenitor populations experience a similar reduction in rRNA transcription but differ in their cellular outcome.

Consistent with broad *tp53* activation, genetic inhibition of *tp53* significantly reduced cell death and improved overall morphology in *polr1c* mutant embryos (Figs. S3, S4), but the effects on the oligodendrocyte lineage were limited. *olig2:egfp* expression showed only modest improvement (Fig. 6) and *mbp:tagRFP* expression was not restored in *polr1c*^-/-^*; tp53^-/-^* zebrafish (Fig. 7). Interestingly, induction of *tp53, cdkn1a*, and *mdm2* were comparable between *polr1c^-/-^; tp53^+/-^*and *polr1c^-/-^; tp53^+/+^* larvae at 3 dpf, indicating that one functional copy of *tp53* is sufficient to trigger a robust transcriptional response to *polr1c* loss. Complete inhibition of *tp53* reduced expression of Tp53 pathway genes to control levels (Fig. 8) but still failed to restore *mbp* expression. Together, these findings suggest that Tp53-dependent cell death contributes to some developmental anomalies caused by *polr1c* deficiency, and impaired myelin development involves additional Tp53-independent mechanisms.

### Developmental timing and requirements for ribosome biogenesis may contribute to tissue-specific responses in *polr1c* mutants

The limited effect of *tp53* inhibition on myelin development in *polr1c* mutants contrasts with its previously described effects on craniofacial development. Tp53-dependent loss of NCC progenitors contributes to craniofacial anomalies in *polr1c* mutant zebrafish and mouse models of TCS, and *tp53* inhibition can ameliorate these craniofacial phenotypes (Jones et al., 2008; Noack Watt et al., 2016). In contrast, preventing Tp53-dependent cell death was not sufficient to restore myelin development in *polr1c* mutants (Fig. 7), suggesting that the consequences of impaired rRNA transcription may depend on the developmental and cellular context.

Developmental timing may contribute to the different responses to *tp53* inhibition. Zebrafish embryos contain maternally supplied rRNAs that persist beyond 2 dpf (Locati et al., 2017) which may transiently support early development despite impaired zygotic rRNA transcription. Consistent with this model, NCCs survive and progress through early developmental stages in *polr1a^-/-^; tp53^-/-^* zebrafish embryos despite a near complete loss of zygotic rRNA transcription, although craniofacial rescue becomes limited later in development (Watt et al., 2018). In *polr1c* mutants, reduced *olig2:egfp* expression emerges by 36 hpf, and myelination initiates at approximately 3 dpf, when there is likely greater reliance on zygotic rRNA transcription. Consequently, preventing apoptosis may preserve early progenitor populations, such as NCCs, but does not correct the defect in rRNA transcription required for ribosome biogenesis and later oligodendrocyte development and myelin production.

Tissue-specific requirements for ribosome biogenesis and translation may also contribute to the phenotypes observed in *polr1c* mutant zebrafish. NCCs are highly proliferative and migratory, requiring high levels of rRNA transcription (Falcon et al., 2022) while ribosome biogenesis initially diminishes during neuronal differentiation (Chau et al., 2018; Ni et al., 2025). Cells with a relatively high rate of rRNA transcription are more susceptible to Tp53-dependent apoptosis following disruption of ribosome biogenesis (Scala et al., 2016). This may contribute to the extensive loss of NCC progenitors in the dorsal neuroepithelium while OPCs activate the Tp53 pathway without undergoing apoptosis. At later stages, the greater requirements for ribosome biogenesis associated with axon growth and myelin production may render their development sensitive to persistent deficits in rRNA transcription even in the absence of cell death. Thus, developmental timing and cell-type-specific requirements may determine how tissues respond to *polr1c* deficiency despite a shared underlying defect in RNA polymerase function.

### rRNA transcription by Pol I is impaired in *polr1c* mutants regardless of *tp53* status

As Polr1c is a shared subunit of Pol I and Pol III, defects in the activity of either polymerase could contribute to the neurological phenotypes observed in *polr1c* mutant zebrafish. Pre-rRNA levels were reduced by approximately 50% in *polr1c* mutant OPCs, remained reduced at the onset of myelination (Figs. 5, 8), and were not improved by *tp53* inhibition (Fig. 8). Thus, inhibition of *tp53* does not restore the underlying defect in Pol I-dependent rRNA transcription. The effects on Pol III-transcribed RNAs were more varied. Some pre-tRNAs were reduced in *polr1c* mutants, while other pre-tRNAs and 7SL RNA were unchanged, and these effects were similarly independent of *tp53* status (Fig. 8). tRNA-sequencing also revealed variable changes in mature tRNA abundance, with either no change or only modest differences at the level of isodecoder (Fig. 8). However, these measurements do not comprehensively assess Pol III function. As mature tRNA abundance reflects a combination of tRNA transcription, processing, and stability, it is not a direct measure of Pol III activity. Additionally, Pol III transcribes a diverse range of non-coding RNAs (White, 2011) beyond the transcripts examined here. Therefore, while we detect robust reduction in Pol I-dependent rRNA transcription, the extent and functional consequences of Pol III dysfunction in this model remain unresolved.

*POLR3*-related leukodystrophy is generally attributed to impaired Pol III activity, and HLD11-associated *POLR1C* variants have been suggested to preferentially disrupt Pol III-mediated transcription (Thiffault et al., 2015). Reduced tRNA production has also been demonstrated in a mouse model of *POLR3*-related leukodystrophy (Moir et al., 2024), supporting an important role for altered Pol III function in disease pathogenesis. In contrast, we recently demonstrated that reduced pre-tRNA transcription in a Pol III-specific zebrafish mutant did not exhibit a corresponding reduction in *mbp:tagRFP* expression at 6 dpf (Lubash et al., 2026). Although myelin structure or later stages of myelination were not assessed, comparison with this model suggests that reduced pre-tRNA transcription alone may not be sufficient to account for the *mbp* phenotype in *polr1c* mutants.

Differences between zebrafish and mammalian models, including developmental timing and maternal contributions, could influence the consequences of Pol III dysfunction on myelination and contribute to the different phenotypes observed across models. Additional effects on other Pol III-transcribed RNAs and impaired ribosome biogenesis may also contribute to impaired myelination in *polr1c* mutants.

The persistent impairment of rRNA transcription in *polr1c* mutant zebrafish could contribute to altered nervous system development, as Pol I dysfunction is also associated with neurological and myelination phenotypes (Kara et al., 2017; Smallwood et al., 2023). Furthermore, myelin formation places substantial demands on protein synthesis. Components of the translational machinery are enriched in CNS myelin transcriptomes (Thakurela et al., 2016), and major myelin proteins such as Mbp rely on local cytoplasmic translation during sheath formation (Colman et al., 1982). Thus, reduced rRNA transcription and ribosome biogenesis could limit the translational capacity required for myelin development even when oligodendrocytes survive and migrate to the appropriate regions in the CNS. Together, these observations support impaired Pol I-dependent rRNA transcription as a contributor to the myelin and neurological phenotypes observed in *polr1c* mutant zebrafish.

There are several limitations to our studies that should be considered. The *polr1c* mutant zebrafish used in our studies represents a global loss of function rather than a specific human disease-associated allele. This broader loss of function may explain differences between our findings and those in variant-specific models. Human *POLR1C* variants may differentially affect Pol I and Pol III assembly or transcription (Misiaszek et al., 2021; Thiffault et al., 2015), and the relative effects on Pol I and Pol III observed in *polr1c* mutant zebrafish may not apply to all *POLR1C*-associated disease variants. In addition, our analysis examined only a small subset of RNAs transcribed by Pol III. A more comprehensive assessment of nascent Pol III-mediated transcription will be required to define how the loss of *polr1c* affects Pol III activity. The limited survival of *polr1c* mutant zebrafish also restricts analysis to early stages of myelin development. Conditional or inducible deletion mammalian models, particularly those targeting *Polr1c* in the oligodendrocyte lineage, could help distinguish cell autonomous requirements for *Polr1c* in later oligodendrocyte development and myelin maintenance. Variant-specific models will likewise be important for defining how individual disease-associated alleles alter Pol I and Pol III function and produce distinct tissue-specific outcomes.

In summary, our study expands the developmental consequences of *polr1c* deficiency to oligodendrocytes and myelination in addition to NCCs. The distinct cellular responses of OPCs and NCCs indicate that a similar stressor can produce distinct outcomes across developing tissues. These distinct outcomes extend to the limited effect of *tp53* inhibition on myelination in *polr1c* mutants. Our findings further identify persistent defects in rRNA transcription by Pol I, and potential contributions from Pol III dysfunction, as potential contributors to nervous system phenotypes in *polr1c* mutants.

Altogether, these findings identify tissue-specific responses to *polr1c* deficiency that may contribute to the pathogenesis of *POLR1C*-associated diseases.

## Materials and Methods

### Ethics statement

All zebrafish work was approved by the Institutional Animal Care and Use Committee at the Stowers Institute for Medical Research (Protocol #2021-124) and the University of Colorado Anschutz Medical Campus (Protocol #01361).

### Zebrafish husbandry and genotyping

Adult zebrafish (*Danio rerio*) between 6-15 months old were maintained on an AB background and housed at 28.5C on a 14-hour light/10-hour dark light cycle at the Stowers Institute Zebrafish Facility and the University of Colorado Anschutz Zebrafish Facility. *polr1c^hi1124^* zebrafish (ZIRC catalog ID# ZL1176) were maintained as heterozygotes, referred to as *polr1c^+/-^* in this manuscript. The *polr1c* wild type allele was detected according to Noack Watt *et al.,* 2016 using the primers forward 5’-CTATTGCTTTTGTCGCATAAAGCG-3’ and reverse 5’-CTCCAGTGTGTTTTCATCTG AAC-3’.

The *polr1c* mutant allele was detected using the primers forward 5’-CTATTGCTTTT GTCGCAT AAAGCG-3’ and reverse 5’-GCTAGCTTGCCAAACCTACAGGT-3’. Adult *polr1c* heterozygous zebrafish were crossed with additional transgenic reporter lines including *Tg(olig2 egfp)* (Shin et al., 2003; gift from B. Appel lab), referred to as *olig2:egfp*, *Tg*(*mbp:<u>tag</u>RFP*) (Hines et al., 2015; gift from B.

Appel lab), referred to as *mbp:tagRFP*, *Tg(sox10:mRFP)* (Kucenas et al., 2008; gift from B. Appel lab) referred to as *sox10:RFP*, and *tp53^m214k^*line, (Berghmans et al., 2005; ZIRC catalog ID# ZL1057), referred to as *tp53^-/-^*. The *tp53* wild-type allele was detected using primers 5’-AGCTGCATGGGGGGGAT-3’ and reverse 5’-GATAGCCTAGTGCGAGCACACTCTT-3’. The *tp53* mutant allele was detected using the primers forward 5’-AGCTGCATGGGGGGGAA-3’ and reverse 5’-GATAGCCTAGTGCGAGCACACTCTT-3’. Zebrafish embryos were raised at 28.5°C and staged according to (Kimmel et al., 1995). When necessary, 1-Phenyl-2-thiourea (0.003%) was added to the embryo media to prevent pigment development.

#### Immunostaining and TUNEL

Whole-mount immunostaining was completed according to Westerfield, 2000 (The Zebrafish Book) using primary antibodies against GFP (1:500, Invitrogen, #A6455; RRID:AB_221570) pHH3 (1:2000, Millipore cat#05-806; RRID:AB_310016), HuC (1:100, Invitrogen cat#A21271; RRID: AB_221448), and Acetylated Tubulin (1:250, Sigma cat#T7451; RRID:AB_609894). Fluorescent secondary antibodies (1:500) included Alexa 488 goat anti-rabbit (Invitrogen Cat# A-11008, RRID:AB_143165), Alexa 546 goat anti-mouse (Invitrogen Cat# A-11030, RRID:AB_144695), and Alexa 647 donkey anti-mouse (Invitrogen Cat# A-31571, RRID:AB_162542). TUNEL was completed with *In Situ* Cell Death Detection Kit, TMR red (Roche Cat# 12156792910). Staining was completed according to (Crump et al., 2004), with slight modifications. TdT/Fluorescein-dUTP reaction was incubated on ice for 1 hour then at 37°C for 1 hour. DAPI staining was conducted at room temperature for 20-30 minutes (1:1000, Invitrogen, Cat#D1306). Embryos were imaged using a 3i Marianas 3D inverted Spinning Disk microscope and a Zeiss LSM 780 camera.

#### Transmission Electron Microscopy

Zebrafish larvae at 5 dpf were genotyped and fixed in EM fixative (2% paraformaldehyde, 2.5% glutaraldehyde, 1% sucrose, 1mM CaCl2, and 50 mM sodium cacodylate (pH 7.4)). After rinsing with rinse buffer (1% sucrose, 1mM CaCl2, and 50 mM sodium cacodylate (pH 7.4)), the tissues were post-fixed with 1% OsO4 and then stained with 0.5% UA overnight. Samples were dehydrated through a graded ethanol series into 100% ethanol, and then with propylene oxide. The samples were infiltrated and embedded in Epon resin (EMS, Fort Washington, PA). EM grids were prepared with a Leica Ultra microtome (Leica UC-6) using diamond knives. The 80 nm sections were post-stained with 4% uranyl acetate in 70% methanol and Sato’s triple lead stain and then imaged with a FEI transmission electron microscope (Tecnai Bio-TWIN 12, FEI).

#### Confocal imaging

For live imaging, embryos were anaesthetized with MS-222 and mounted in 0.8% low melting agarose in E2 media. For timelapse imaging, images were captured every 12 minutes for 15 hours with a 40x objective on a Zeiss 780 microscope fitted with an incubation chamber kept at 28°C. Videos were processed using ImageJ. To image *mbp:tagRFP* and *olig2:egfp* expression, larvae were mounted in 0.8% low melting point agarose with MS-222 and imaged on a Zeiss 700 confocal, Zeiss 780 confocal, or Andor Dragonfly spinning disk confocal with a 20x objective. For imaging of fixed samples, embryos were mounted in 0.8% low melting agarose in PBS and imaged on the Zeiss 700 confocal or 3i Marianas 3D inverted Spinning Disk with a 10x or 20x objective.

#### Image analysis

Images were analyzed utilizing Imaris 9. Regions of interest were defined based on anatomical landmarks, and the region was the same size across all images analyzed. Surfaces were generated using automated settings in Imaris and volumes were recorded in µm^3^. Cell counts were determined within the region of interest using the spots tool. Results were analyzed for statistical significance using GraphPad Prism 10.

#### Image processing

Figures were assembled in Adobe Photoshop. Any adjustments to image brightness and contrast were applied equally across the entire image.

#### Cell dissociation, cell sorting, and RNA extraction

Embryos were screened for expression of *olig2:egfp* and *sox10:mRFP* and then manually dechorionated. Samples that were *olig2:egfp+* only, *sox10:mRFP+* only, and without transgenic reporter expression were prepared for use as gating controls in parallel with the *polr1c* control and mutant samples. 25 *olig2:egfp+; sox10:mRFP+* embryos per sample were de-yolked with calcium-free Ringer’s solution and washed with dPBS. Samples were centrifuged at 350 x *g* at room temperature and the supernatant was removed. TypLE (Gibco) was added to each sample and incubated at 29°C for 15 minutes and quenched with 100% fetal bovine serum. Cells were spun down at 350 x *g* at 4°C, washed with dPBS, centrifuged again, and the supernatant was removed. Cell pellets were resuspended in dPBS and filtered through a 30 µm filter. Prior to analysis, cells were stained with DAPI (Sigma Chemicals) at 2 µL/mL to identify cells with compromised membrane integrity (dying/dead cells), and with DRAQ5 (BioStatus Limited) at 25 µM final concentration to label nucleated cells. Flow cytometric analysis and fluorescence-activated cell sorting (FACS) were performed on a BD Influx cell sorter (BD Biosciences) equipped with a 100 µm nozzle tip and operated at a sheath pressure of 20 PSI. Sheath fluid consisted of particulate-free PBS supplemented with 0.1% Pluronic F-68. Cells were gated sequentially as follows: (1) a broad forward scatter (FSC) / side scatter (SSC) gate excluding debris and the highest few percent of events by scatter intensity; (2) an FSC-based doublet discrimination gate to exclude non-singlet events; and (3) a rectangular gate identifying a clear and reproducible DRAQ5-positive, DAPI-negative population, hereafter referred to as Live Single Cells. Live Single Cells were subsequently gated and sorted based on GFP and RFP fluorescence intensity. Post-sort purity was verified by immediate reanalysis of sorted fractions on the same instrument. Representative reanalyzes demonstrated post-sort purities of ≥92%, with the primary contaminating events consistent with dying cells and sub-cellular debris. Following sorting, cells were pelleted by centrifugation and then immediately lysed in TRIzol. RNA was extracted using the Qiagen miRNA micro kit and concentrations were measured on the Nanodrop 2000. The SuperScript III kit (Invitrogen, cat#18080044) was used for cDNA synthesis and then used in qRT-PCR (described below).

#### RNA extraction and cDNA synthesis

Samples were collected for RNA extraction at 24 hpf and 3 dpf. Heads were placed in RNAlater solution (Invitrogen, AM7020) while tails were lysed for genotyping.

Embryos of the same genotype from the same clutch were pooled in a single RNA sample, with a minimum of 3 heads per sample. For the 24 hpf samples, RNA was extracted using the Qiagen miRNeasy Micro Kit with on-column DNase treatment. The concentration and purity were determined on the Nanodrop 1000 Spectrophotometer. For samples collected at 3 dpf, heads of the same genotype were pooled and lysed in TRIzol followed purification and on-column DNase treatment using the Zymo Clean & Concentrator kit. Equal amounts of RNA were used to synthesis cDNA for qPCR with the Superscript III kit (Invitrogen, Cat#18080044) or Superscript IV kit (Invitrogen, Cat#18090010) using random hexamers.

#### qRT-PCR

After cDNA synthesis, cDNA was diluted to the required concentration determined by standard curve analysis for each assay. PerfeCTa (Quanta Biosciences) reaction mix and the ABI 7900HT real time PCR cycler or the iTaq Sybr reaction mix (BioRad) and the BioRad CFX real time PCR cycler were used measure cDNA amplification. Three biological replicates were run in technical triplicate for each experiment. No template and no reverse-transcriptase samples were run as negative controls. Relative quantification was determined using the 2^-ΔΔCT^ method (Livak and Schmittgen, 2001).

Primers used for qRT-PCR are listed in Table 1.

**Table 1.**
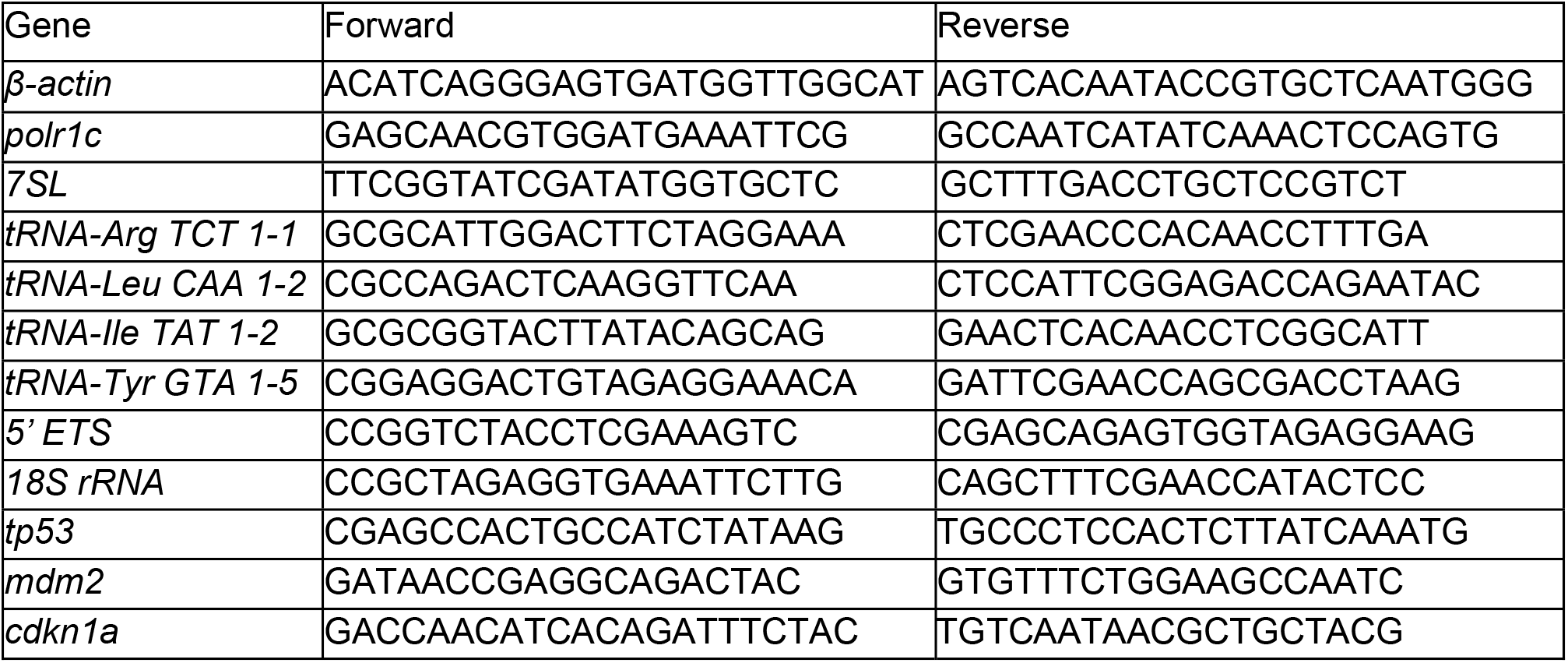
Primer sequences for quantitative RT-PCR experiments.

#### RNA-sequencing

RNA was collected from 24 hpf zebrafish embryos as described above and tested for quality on an Agilent 2100 Bioanalyzer. Three samples of five embryos per sample were collected for both wild-types and mutants. Samples with an RNA integrity (RIN) score greater than 8.0 were used for library construction. RNA underwent poly(A) selection prior to library construction and sequencing. Single end 50 base sequence reads generated on a HiSeq instrument from Illumina. Reads were mapped to the Danio rerio genome version GRCz11 with gene descriptions from Ensembl 115 by STAR version 2.7.11b using --quantMode GeneCounts and default parameters. Read mappings were assessed for differential expression between conditions using R and Bioconductor with edgeR (Robinson et al., 2010). The resulting p-values were adjusted by the method of Benjamini and Hochberg (Benjamini and Hochberg, 1995). For pathway analysis, we utilized software from the Bioinformatics and Biostatistics Share Resource at the University of Colorado Anschutz (bioinformatics.cuanschutz.edu). The input gene lists for Gene Ontology analysis had an adjusted p-value of at least 0.05 and fold change of at least 2.

#### tRNA-sequencing

Total RNA was collected from *polr1c* wild-type and homozygous mutant heads at 3 dpf. The purity and concentration were determined using the NanoDrop and tRNA-sequencing service was provided by Arraystar, Inc. Briefly, tRNAs were purified from 4µg total RNA, m1A and m3C demethylated using the rtStar*TM* tRF&tiRNA Pretreatment Kit, and partially hydrolyzed according to the Hydro-tRNAseq method (Gogakos et al., 2017). Fragments of ∼19-35 nucleotides for partially hydrolyzed and re-phosphorylated tRNAs were prepared for small RNA sequencing using the NEBNext Multiplex Small RNA Library Prep set of Illumina kit. The tRNA-seq libraries were quantified using Agilent 2100 BioAnalyzer and then equally mixed and sequenced on an Illumina Nextseq 550 according to the manufacturer’s instructions.

#### tRNA-seq analysis

Sequence quality was examined by FastQC software and trimmed reads (trimmed 3’-adaptor bases by cutadapt) were aligned to the cytoplasmic mature-tRNA sequences from GtRNAdb and the mitochondrial tRNA sequences from mitotRNAdb using the *Danio rerio* genome version GRCz11 and BWA software (Li and Durbin, 2009). For tRNA alignment, the maximum mismatch was 2 (Gogakos et al., 2017). Expression profiling of tRNAs was calculated based on uniquely mapped reads and including multi mapped reads, respectively. The differentially expressed tRNAs were screened based on the count value with R package edgeR (Robinson et al., 2010).

### Statistics

A two-tailed Student’s t-test was used to determine significance between the means with p<0.05 deemed statistically significant. qPCR data was analyzed using GraphPad Prism v.11.0.2. For analysis of image quantification, one-way ANOVA was performed to determine if there were significant differences between groups. If the ANOVA results were significant, the Tukey test was used to determine which differences were significant, or the Tukey-Kramer was used when groups had a differing number of samples.

## Acknowledgements

The authors thank Dr. Bruce Appel for reagents and advice; the staff in the Stowers Institute for Medical Research zebrafish facility and the staff in the University of Colorado Anschutz zebrafish facility for zebrafish care and maintenance; the staff in the Stowers Institute for Medical Research Flow Cytometry core for cell sorting assistance; and the personnel from the University of Colorado Anschutz Advanced Light Microscopy Core for imaging assistance. Imaging at the University of Colorado Anschutz was performed in the Advanced Light Microscopy Core facility of the NeuroTechnology Center, which is supported in part by Rocky Mountain Neurological Disorders Core Grant (P30NS048154) and by Diabetes Research Center Grant (P30 DK116073). This study was partly supported by the National Institutes of Health P30CA046934 by utilizing the tools created by the Bioinformatics and Biostatistics Shared Resource.

## Competing interests

The authors declare no competing interests.

## Funding

This work was supported by the National Institute of Dental and Craniofacial Research [F31 DE023017 to K.E.N.W.; K99/R00 DE030971 to K.E.N.W.], and funds from the University of Colorado Anschutz School of Dental Medicine. The content is solely the responsibility of the authors and does not necessarily represent the official views of the NIH.

## Data and resource availability

All details and resources are available within the article and supplementary information. Raw sequencing data will be available upon publication from a publicly available database.

## Supporting Information

**Fig. S1. Myelin is reduced in *polr1c^-/-^* zebrafish at 7 dpf.** A-D) Expression of *Tg(olig2:egfp)* and *Tg(mbp:tagRFP)* in control and *polr1c^-/-^* zebrafish reveals a reduced percentage of *olig2+; mbp+* cells (C) and a significant reduction in overall *mbp:tagRFP* volume in mutant zebrafish (D). n = 13 controls, 12 mutants. E-H) Electron microscopy was used to assess myelin structure in the spinal cord at 7 dpf. Compacted myelin around axons was detected in the central nervous system in controls and *polr1c^-/-^* zebrafish. Strikingly, this was not the case in the neural crest cell-derived peripheral nervous system.

**Fig. S2. Staining for HuC and Isl1 reveal slight changes in neuronal populations at 36 hpf in *polr1c^-/-^*embryos. A,B)** Immunofluorescence staining for pan-neuronal marker HuC appears similar between controls and *polr1c^-/-^* zebrafish at 36 hpf. **C-F)** Immunofluorescence staining with Isl1 was used to assess motor neurons (C,D). While no significant differences were detected in the head (E), a significant reduction in staining was detected in the trunk (F). n = 7 controls, 8 mutants. Scale bar = 100 µm

**Fig. S3.** Single channel images of pHH3 (white), *olig2:egfp* (green), and TUNEL (magenta) at 26 hpf from embryos represented in Fig. 6. Scale bar = 100 µm

**Fig. S4.** Single channel images of pHH3 (white), *olig2:egfp* (green), and TUNEL (magenta) at 36 hpf from embryos represented in Fig. 6. Scale bar = 100 µm

**Fig. S5.** Single channel images of pHH3 (white), *olig2:egfp* (green), and TUNEL (magenta) at 72 hpf from embryos represented in Fig. 6. Scale bar = 100 µm

**Fig. S6.** Proliferation is not significantly different in *polr1c^-/-^; tp53^-^* zebrafish compared to controls.

## Notes

### Competing Interest Statement

The authors have declared no competing interest.

## References

Ackerman, S. D. and Monk, K. R. (2016). The scales and tales of myelination: using zebrafish and mouse to study myelinating glia. Brain Research 1641, 79–91.

Bechler, Marie E., Byrne, L. and ffrench-Constant, C. (2015). CNS Myelin Sheath Lengths Are an Intrinsic Property of Oligodendrocytes. Current Biology 25, 2411–2416.

Benjamini, Y. and Hochberg, Y. (1995). Controlling the False Discovery Rate: A Practical and Powerful Approach to Multiple Testing. Journal of the Royal Statistical Society: Series B (Methodological) 57, 289–300.

Berghmans, S., Murphey, R. D., Wienholds, E., Neuberg, D., Kutok, J. L., Fletcher, C. D., Morris, J. P., Liu, T. X., Schulte-Merker, S., Kanki, J. P. et al. (2005). tp53 mutant zebrafish develop malignant peripheral nerve sheath tumors. Proc Natl Acad Sci U S A 102, 407–12.

Bhatt, S., Diaz, R. and Trainor, P. A. (2013). Signals and switches in Mammalian neural crest cell differentiation. Cold Spring Harb Perspect Biol 5.

Boguta, M. (2022). Assembly of RNA polymerase III complex involves a putative co-translational mechanism. Gene 824, 146394.

Chau, K. F., Shannon, M. L., Fame, R. M., Fonseca, E., Mullan, H., Johnson, M. B., Sendamarai, A. K., Springel, M. W., Laurent, B. and Lehtinen, M. K. (2018). Downregulation of ribosome biogenesis during early forebrain development. eLife 7, e36998.

Colman, D. R., Kreibich, G., Frey, A. B. and Sabatini, D. D. (1982). Synthesis and incorporation of myelin polypeptides into CNS myelin. J Cell Biol 95, 598–608.

Crump, J. G., Swartz, M. E. and Kimmel, C. B. (2004). An integrin-dependent role of pouch endoderm in hyoid cartilage development. PLoS Biol 2, E244.

Dauwerse, J. G., Dixon, J., Seland, S., Ruivenkamp, C. A., van Haeringen, A., Hoefsloot, L. H., Peters, D. J., Boers, A. C., Daumer-Haas, C., Maiwald, R. et al. (2011). Mutations in genes encoding subunits of RNA polymerases I and III cause Treacher Collins syndrome. Nat Genet 43, 20–2.

de Monasterio-Schrader, P., Jahn, O., Tenzer, S., Wichert, S. P., Patzig, J. and Werner, H. B. (2012). Systematic approaches to central nervous system myelin. Cellular and Molecular Life Sciences 69, 2879–2894.

DeMyer, W., Zeman, W. and Palmer, C. G. (1964). THE FACE PREDICTS THE BRAIN: DIAGNOSTIC SIGNIFICANCE OF MEDIAN FACIAL ANOMALIES FOR HOLOPROSENCEPHALY (ARHINENCEPHALY). Pediatrics 34, 256–263.

Donati, G., Peddigari, S., Mercer, C. A. and Thomas, G. (2013). 5S ribosomal RNA is an essential component of a nascent ribosomal precursor complex that regulates the Hdm2-p53 checkpoint. Cell Rep 4, 87–98.

Dutt, S., Narla, A., Lin, K., Mullally, A., Abayasekara, N., Megerdichian, C., Wilson, F. H., Currie, T., Khanna-Gupta, A., Berliner, N. et al. (2011). Haploinsufficiency for ribosomal protein genes causes selective activation of p53 in human erythroid progenitor cells. Blood 117, 2567–76.

Falcon, K. T., Watt, K. E. N., Dash, S., Zhao, R., Sakai, D., Moore, E. L., Fitriasari, S., Childers, M., Sardiu, M. E., Swanson, S. et al. (2022). Dynamic regulation and requirement for ribosomal RNA transcription during mammalian development. Proceedings of the National Academy of Sciences 119, e2116974119.

Gauquelin, L., Cayami, F. K., Sztriha, L., Yoon, G., Tran, L. T., Guerrero, K., Hocke, F., van Spaendonk, R. M. L., Fung, E. L., D’Arrigo, S. et al. (2019). Clinical spectrum of POLR3-related leukodystrophy caused by biallelic POLR1C pathogenic variants. Neurol Genet 5, e369.

Girbig, M., Misiaszek, A. D., Vorländer, M. K., Lafita, A., Grötsch, H., Baudin, F., Bateman, A. and Müller, C. W. (2021). Cryo-EM structures of human RNA polymerase III in its unbound and transcribing states. Nature Structural & Molecular Biology 28, 210–219.

Gogakos, T., Brown, M., Garzia, A., Meyer, C., Hafner, M. and Tuschl, T. (2017). Characterizing Expression and Processing of Precursor and Mature Human tRNAs by Hydro-tRNAseq and PAR-CLIP. Cell Rep 20, 1463–1475.

Grummt, I. (2003). Life on a planet of its own: regulation of RNA polymerase I transcription in the nucleolus. Genes & Development 17, 1691–1702.

Haupt, Y., Maya, R., Kazaz, A. and Oren, M. (1997). Mdm2 promotes the rapid degradation of p53. Nature 387, 296–299.

Hines, J. H., Ravanelli, A. M., Schwindt, R., Scott, E. K. and Appel, B. (2015). Neuronal activity biases axon selection for myelination in vivo. Nat Neurosci 18, 683–689.

Honda, R., Tanaka, H. and Yasuda, H. (1997). Oncoprotein MDM2 is a ubiquitin ligase E3 for tumor suppressor p53. FEBS Letters 420, 25–27.

Inoue, A., Takahashi, M., Hatta, K., Hotta, Y. and Okamoto, H. (1994). Developmental regulation of Islet-1 mRNA expression during neuronal differentiation in embryonic zebrafish. Developmental Dynamics 199, 1–11.

Jones, N. C., Lynn, M. L., Gaudenz, K., Sakai, D., Aoto, K., Rey, J. P., Glynn, E. F., Ellington, L., Du, C., Dixon, J. et al. (2008). Prevention of the neurocristopathy Treacher Collins syndrome through inhibition of p53 function. Nat Med 14, 125–33.

Kalita, K., Makonchuk, D., Gomes, C., Zheng, J. J. and Hetman, M. (2008). Inhibition of nucleolar transcription as a trigger for neuronal apoptosis. J Neurochem 105, 2286–99.

Kara, B., Köroğlu, Ç., Peltonen, K., Steinberg, R. C., Maraş Genç, H., Hölttä-Vuori, M., Güven, A., Kanerva, K., Kotil, T., Solakoğlu, S. et al. (2017). Severe neurodegenerative disease in brothers with homozygous mutation in POLR1A. Eur J Hum Genet 25, 315–323.

Kim, C.-H., Ueshima, E., Muraoka, O., Tanaka, H., Yeo, S.-Y., Huh, T.-L. and Miki, N. (1996). Zebrafish elav/HuC homologue as a very early neuronal marker. Neuroscience Letters 216, 109–112.

Kimmel, C. B., Ballard, W. W., Kimmel, S. R., Ullmann, B. and Schilling, T. F. (1995). Stages of embryonic development of the zebrafish. Dev Dyn 203, 253–310.

Knecht, A. K. and Bronner-Fraser, M. (2002). Induction of the neural crest: a multigene process. Nat Rev Genet 3, 453–61.

Korzh, V., Edlund, T. and Thor, S. (1993). Zebrafish primary neurons initiate expression of the LIM homeodomain protein Isl-1 at the end of gastrulation. Development 118, 417–425.

Kubbutat, M. H. G., Jones, S. N. and Vousden, K. H. (1997). Regulation of p53 stability by Mdm2. Nature 387, 299–303.

Kucenas, S., Takada, N., Park, H.-C., Woodruff, E., Broadie, K. and Appel, B. (2008). CNS-derived glia ensheath peripheral nerves and mediate motor root development. Nature Neuroscience 11, 143–151.

Laferté, A., Favry, E., Sentenac, A., Riva, M., Carles, C. and Chédin, S. (2006). The transcriptional activity of RNA polymerase I is a key determinant for the level of all ribosome components. Genes Dev 20, 2030–2040.

Lafontaine, D. L. J. and Tollervey, D. (2001). The function and synthesis of ribosomes. Nature Reviews Molecular Cell Biology 2, 514–520.

Lata, E., Choquet, K., Sagliocco, F., Brais, B., Bernard, G. and Teichmann, M. (2021). RNA Polymerase III Subunit Mutations in Genetic Diseases. Front Mol Biosci Volume 8–2021.

Li, H. and Durbin, R. (2009). Fast and accurate short read alignment with Burrows-Wheeler transform. Bioinformatics 25, 1754–60.

Livak, K. J. and Schmittgen, T. D. (2001). Analysis of Relative Gene Expression Data Using Real-Time Quantitative PCR and the 2−ΔΔCT Method. Methods 25, 402–408.

Locati, M. D., Pagano, J. F. B., Girard, G., Ensink, W. A., van Olst, M., van Leeuwen, S., Nehrdich, U., Spaink, H. P., Rauwerda, H., Jonker, M. J. et al. (2017). Expression of distinct maternal and somatic 5.8S, 18S, and 28S rRNA types during zebrafish development. Rna 23, 1188–1199.

Löhr, K., Möritz, C., Contente, A. and Dobbelstein, M. (2003). p21/CDKN1A mediates negative regulation of transcription by p53. J Biol Chem 278, 32507–16.

Lubash, B. T., Gutierrez, R., Hansen, N. A., Fink, K., Hopkins, C. A., Sands, L. B., Nelson, J. C. and Watt, K. E. N. (2026). RNA Polymerase III subunit Polr3a is required for craniofacial cartilage and bone development in zebrafish. PLOS Genet 22, e1012164.

Macintosh, J., Michell-Robinson, M., Chen, X. and Bernard, G. (2023). Decreased RNA polymerase III subunit expression leads to defects in oligodendrocyte development. Frontiers in Neuroscience Volume 17–2023.

McGowan, K. A., Li, J. Z., Park, C. Y., Beaudry, V., Tabor, H. K., Sabnis, A. J., Zhang, W., Fuchs, H., de Angelis, M. H., Myers, R. M. et al. (2008). Ribosomal mutations cause p53-mediated dark skin and pleiotropic effects. Nat Genet 40, 963–70.

Mirchi, A., Guay, S.-P., Tran, L. T., Wolf, N. I., Vanderver, A., Brais, B., Sylvain, M., Pohl, D., Rossignol, E., Saito, M. et al. (2023). Craniofacial features of POLR3-related leukodystrophy caused by biallelic variants in POLR3A, POLR3B and POLR1C. Journal of Medical Genetics 60, 1026.

Misiaszek, A. D., Girbig, M., Grötsch, H., Baudin, F., Murciano, B., Lafita, A. and Müller, C. W. (2021). Cryo-EM structures of human RNA polymerase I. Nature Structural & Molecular Biology 28, 997–1008.

Moir, R. D., Merheb, E., Chitu, V., Stanley, E. R. and Willis, I. M. (2024). Molecular basis of neurodegeneration in a mouse model of Polr3-related disease. eLife 13, RP95314.

Moir, R. D. and Willis, I. M. (2013). Regulation of pol III transcription by nutrient and stress signaling pathways. Biochimica et Biophysica Acta (BBA) - Gene Regulatory Mechanisms 1829, 361–375.

Murray, J. A. and Blakemore, W. F. (1980). The relationship between internodal length and fibre diameter in the spinal cord of the cat. Journal of the Neurological Sciences 45, 29–41.

Ni, C. and Buszczak, M. (2023). The homeostatic regulation of ribosome biogenesis. Seminars in Cell & Developmental Biology 136, 13–26.

Ni, C., Wei, Y., Vona, B., Park, D., Wei, Y., Schmitz, D. A., Ding, Y., Sakurai, M., Ballard, E., Li, L. et al. (2025). A programmed decline in ribosome levels governs human early neurodevelopment. Nature Cell Biology 27, 1240–1255.

Noack Watt, K. E., Achilleos, A., Neben, C. L., Merrill, A. E. and Trainor, P. A. (2016). The Roles of RNA Polymerase I and III Subunits Polr1c and Polr1d in Craniofacial Development and in Zebrafish Models of Treacher Collins Syndrome. PLoS Genet 12, e1006187.

Park, H.-C., Mehta, A., Richardson, J. S. and Appel, B. (2002). olig2 Is Required for Zebrafish Primary Motor Neuron and Oligodendrocyte Development. Developmental Biology 248, 356–368.

Parlato, R., Kreiner, G., Erdmann, G., Rieker, C., Stotz, S., Savenkova, E., Berger, S., Grummt, I. and Schütz, G. (2008). Activation of an Endogenous Suicide Response after Perturbation of rRNA Synthesis Leads to Neurodegeneration in Mice. The Journal of Neuroscience 28, 12759.

Ramsay, E. P., Abascal-Palacios, G., Daiß, J. L., King, H., Gouge, J., Pilsl, M., Beuron, F., Morris, E., Gunkel, P., Engel, C. et al. (2020). Structure of human RNA polymerase III. Nat Commun 11, 6409.

Ravanelli, A. M. and Appel, B. (2015). Motor neurons and oligodendrocytes arise from distinct cell lineages by progenitor recruitment. Genes & Development 29, 2504–2515.

Robinson, M. D., McCarthy, D. J. and Smyth, G. K. (2010). edgeR: a Bioconductor package for differential expression analysis of digital gene expression data. Bioinformatics 26, 139–40.

Rubbi, C. P. and Milner, J. (2003). Disruption of the nucleolus mediates stabilization of p53 in response to DNA damage and other stresses. EMBO J 22, 6068–77.

Sakai, D., Dixon, J., Achilleos, A., Dixon, M. and Trainor, P. A. (2016). Prevention of Treacher Collins syndrome craniofacial anomalies in mouse models via maternal antioxidant supplementation. Nat Commun 7, 10328.

Scala, F., Brighenti, E., Govoni, M., Imbrogno, E., Fornari, F., Treré, D., Montanaro, L. and Derenzini, M. (2016). Direct relationship between the level of p53 stabilization induced by rRNA synthesis-inhibiting drugs and the cell ribosome biogenesis rate. Oncogene 35, 977–989.

Shin, J., Park, H.-C., Topczewska, J. M., Mawdsley, D. J. and Appel, B. (2003). Neural cell fate analysis in zebrafish using olig2 BAC transgenics. Methods in Cell Science 25, 7–14.

Simmons, T. and Appel, B. (2012). Mutation of pescadillo Disrupts Oligodendrocyte Formation in Zebrafish. PLoS One 7, e32317.

Sloan, K. E., Bohnsack, M. T. and Watkins, N. J. (2013). The 5S RNP couples p53 homeostasis to ribosome biogenesis and nucleolar stress. Cell Rep 5, 237–47.

Smallwood, K., Watt, K. E. N., Ide, S., Baltrunaite, K., Brunswick, C., Inskeep, K., Capannari, C., Adam, M. P., Begtrup, A., Bertola, D. R. et al. (2023). POLR1A variants underlie phenotypic heterogeneity in craniofacial, neural, and cardiac anomalies. The American Journal of Human Genetics 110, 809–825.

Thakurela, S., Garding, A., Jung, R. B., Müller, C., Goebbels, S., White, R., Werner, H. B. and Tiwari, V. K. (2016). The transcriptome of mouse central nervous system myelin. Scientific Reports 6, 25828.

Thiffault, I., Wolf, N. I., Forget, D., Guerrero, K., Tran, L. T., Choquet, K., Lavallée-Adam, M., Poitras, C., Brais, B., Yoon, G. et al. (2015). Recessive mutations in POLR1C cause a leukodystrophy by impairing biogenesis of RNA polymerase III. Nat Commun 6, 7623.

Walker-Kopp, N., Jackobel, A. J., Pannafino, G. N., Morocho, P. A., Xu, X. and Knutson, B. A. (2017). Treacher Collins syndrome mutations in Saccharomyces cerevisiae destabilize RNA polymerase I and III complex integrity. Hum Mol Genet 26, 4290–4300.

Watt, K. E. N., Macintosh, J., Bernard, G. and Trainor, P. A. (2023). RNA Polymerases I and III in development and disease. Seminars in Cell & Developmental Biology 136, 49–63.

Watt, K. E. N., Neben, C. L., Hall, S., Merrill, A. E. and Trainor, P. A. (2018). tp53-dependent and independent signaling underlies the pathogenesis and possible prevention of Acrofacial Dysostosis–Cincinnati type. Human Molecular Genetics 27, 2628–2643.

Weaver, K. N., Watt, K. E., Hufnagel, R. B., Navajas Acedo, J., Linscott, L. L., Sund, K. L., Bender, P. L., König, R., Lourenco, C. M., Hehr, U. et al. (2015). Acrofacial Dysostosis, Cincinnati Type, a Mandibulofacial Dysostosis Syndrome with Limb Anomalies, Is Caused by POLR1A Dysfunction. Am J Hum Genet 96, 765–74.

White, R. J. (2011). Transcription by RNA polymerase III: more complex than we thought. Nature Reviews Genetics 12, 459–463.

Wild, T. and Cramer, P. (2012). Biogenesis of multisubunit RNA polymerases. Trends in Biochemical Sciences 37, 99–105.

Wilson, S. W., Ross, L. S., Parrett, T. and Easter, S. S., Jr. (1990). The development of a simple scaffold of axon tracts in the brain of the embryonic zebrafish, Brachydanio rerio. Development 108, 121–145.

Woolnough, J. L., Atwood, B. L., Liu, Z., Zhao, R. and Giles, K. E. (2016). The Regulation of rRNA Gene Transcription during Directed Differentiation of Human Embryonic Stem Cells. PLoS One 11, e0157276.

Xia, W. and Fancy, S. P. J. (2021). Mechanisms of oligodendrocyte progenitor developmental migration. Developmental Neurobiology 81, 985–996.

Yelick, P. C. and Trainor, P. A. (2015). Ribosomopathies: Global process, tissue specific defects. Rare Dis 3, e1025185.

Zannino, D. A. and Appel, B. (2009). Olig2^+^ Precursors Produce Abducens Motor Neurons and Oligodendrocytes in the Zebrafish Hindbrain. The Journal of Neuroscience 29, 2322–2333.

Zhang, Q., Shalaby, N. A. and Buszczak, M. (2014). Changes in rRNA transcription influence proliferation and cell fate within a stem cell lineage. Science 343, 298–301.

