## Supplementary figures and images for "Loss of *polr1c* disrupts myelination in a zebrafish model of *POLR1C*-associated disease"

### Fig. S1

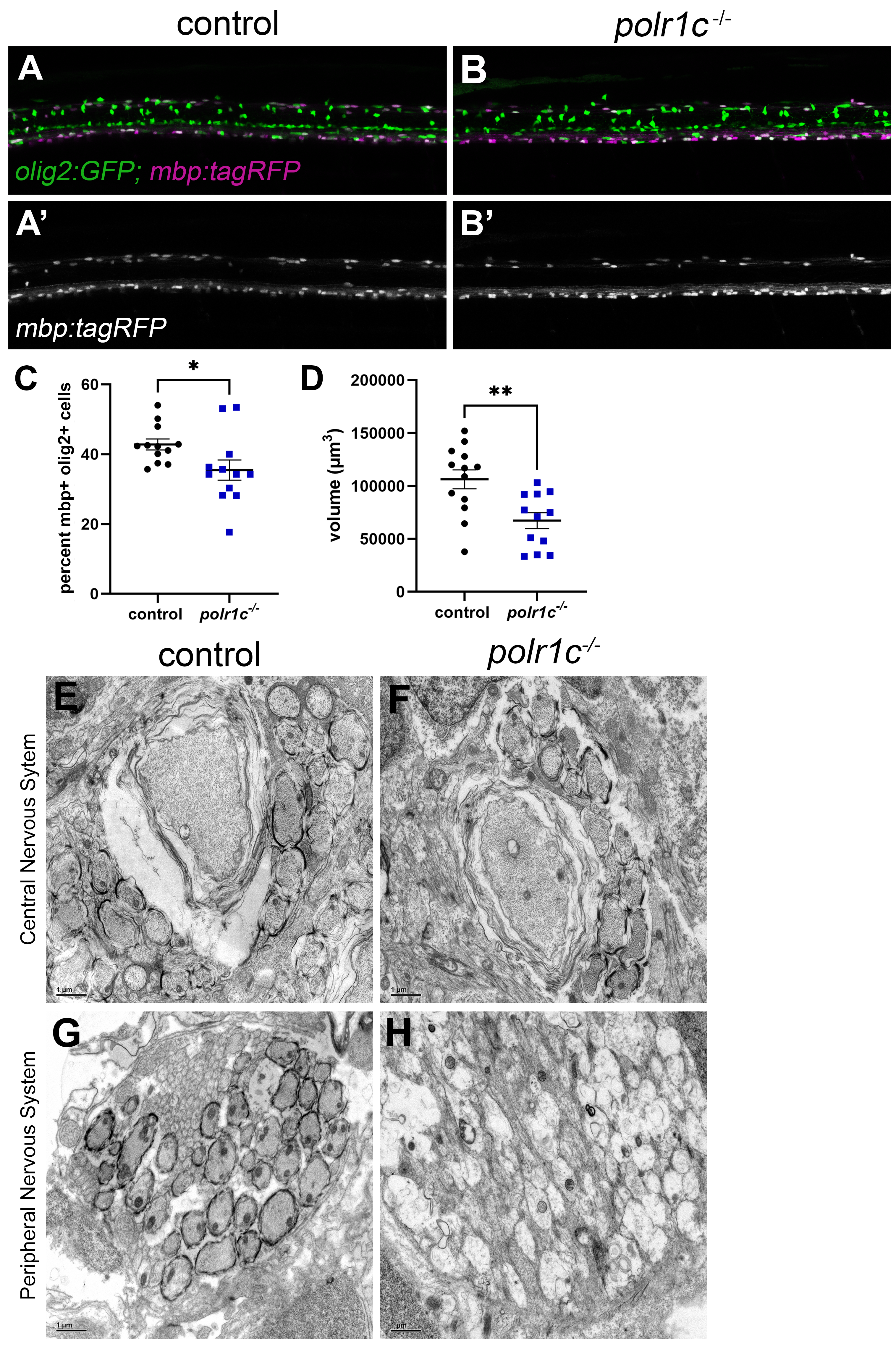

### Fig. S2

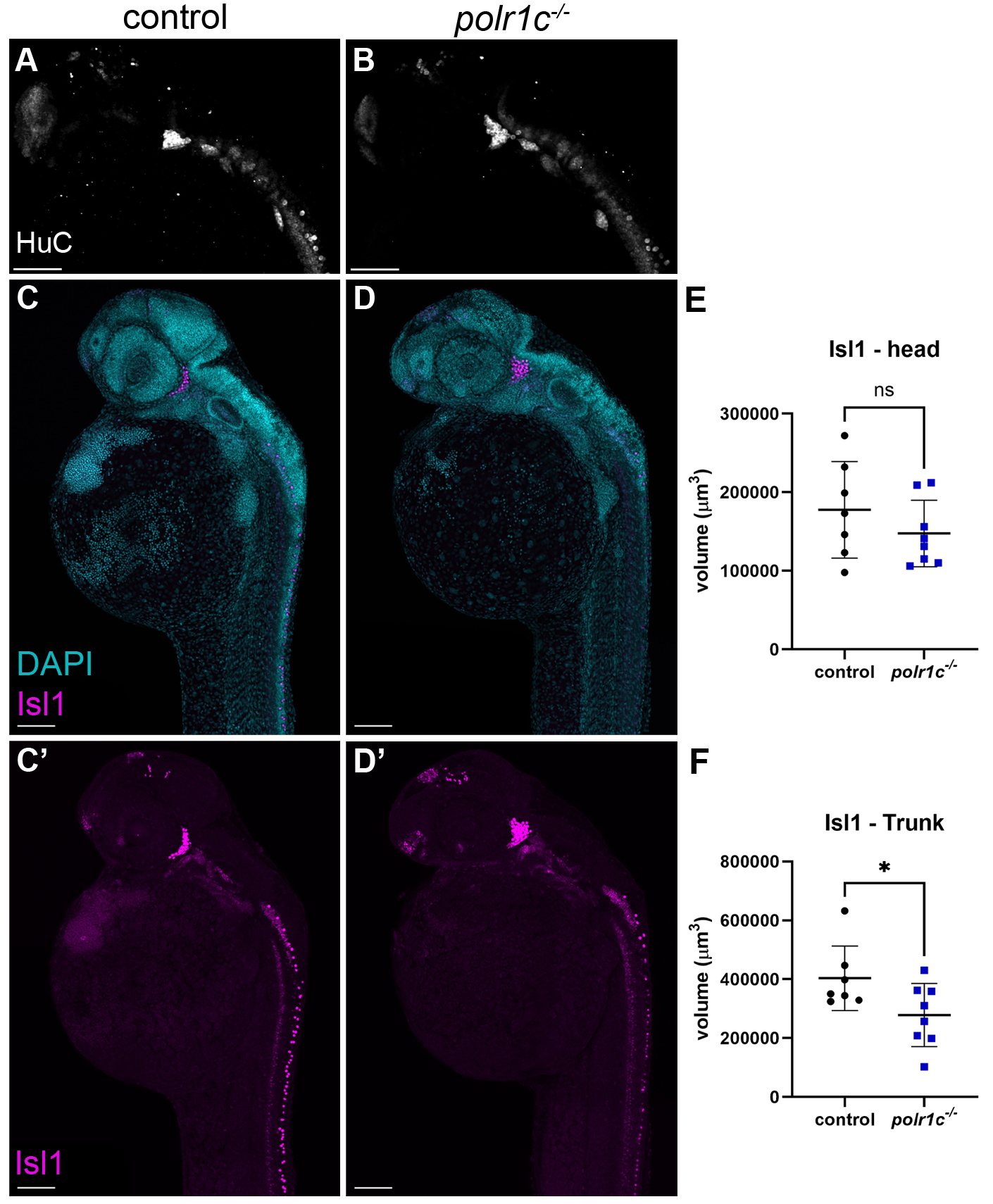

### Fig. S3

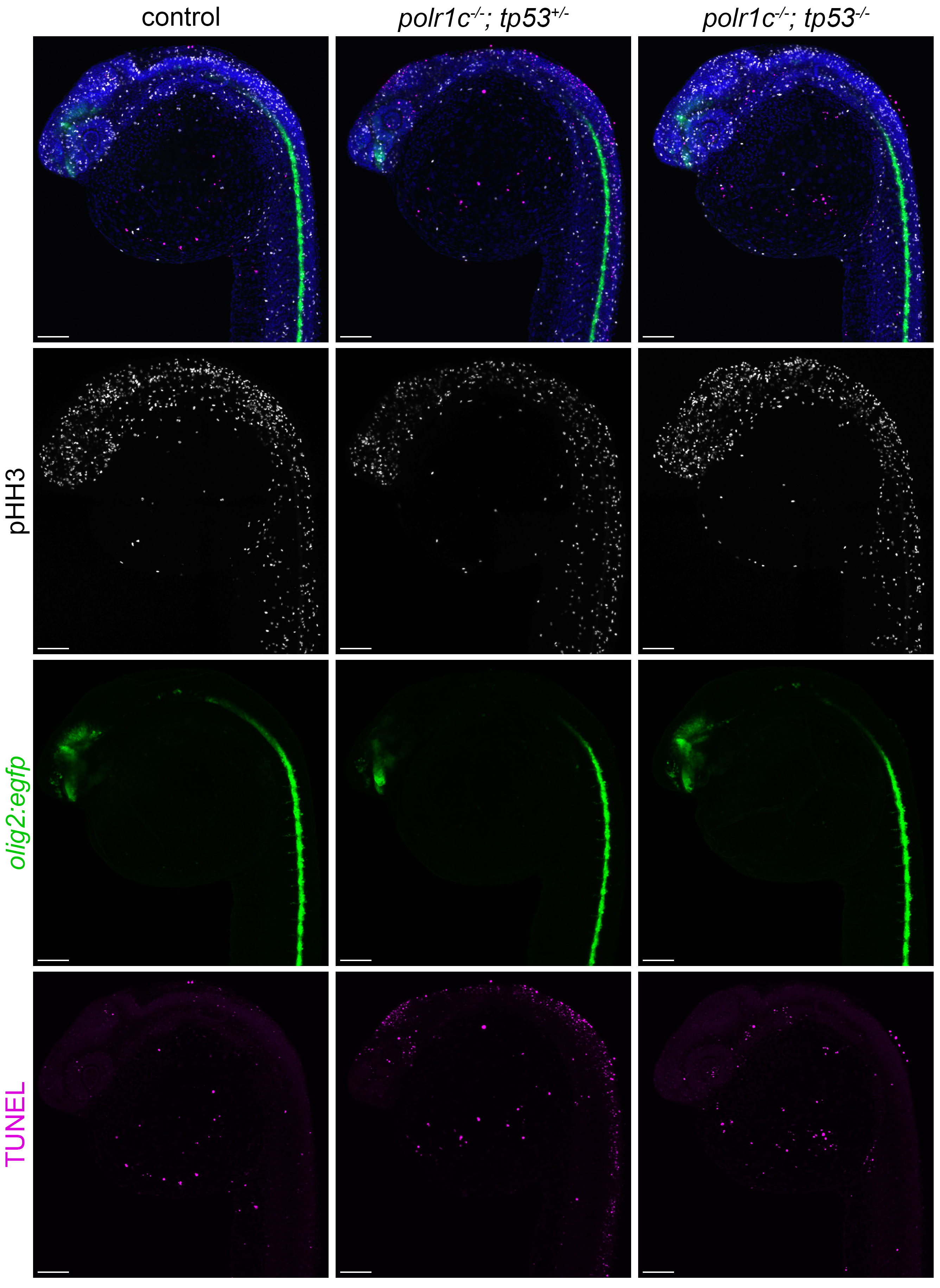

### Fig. S4

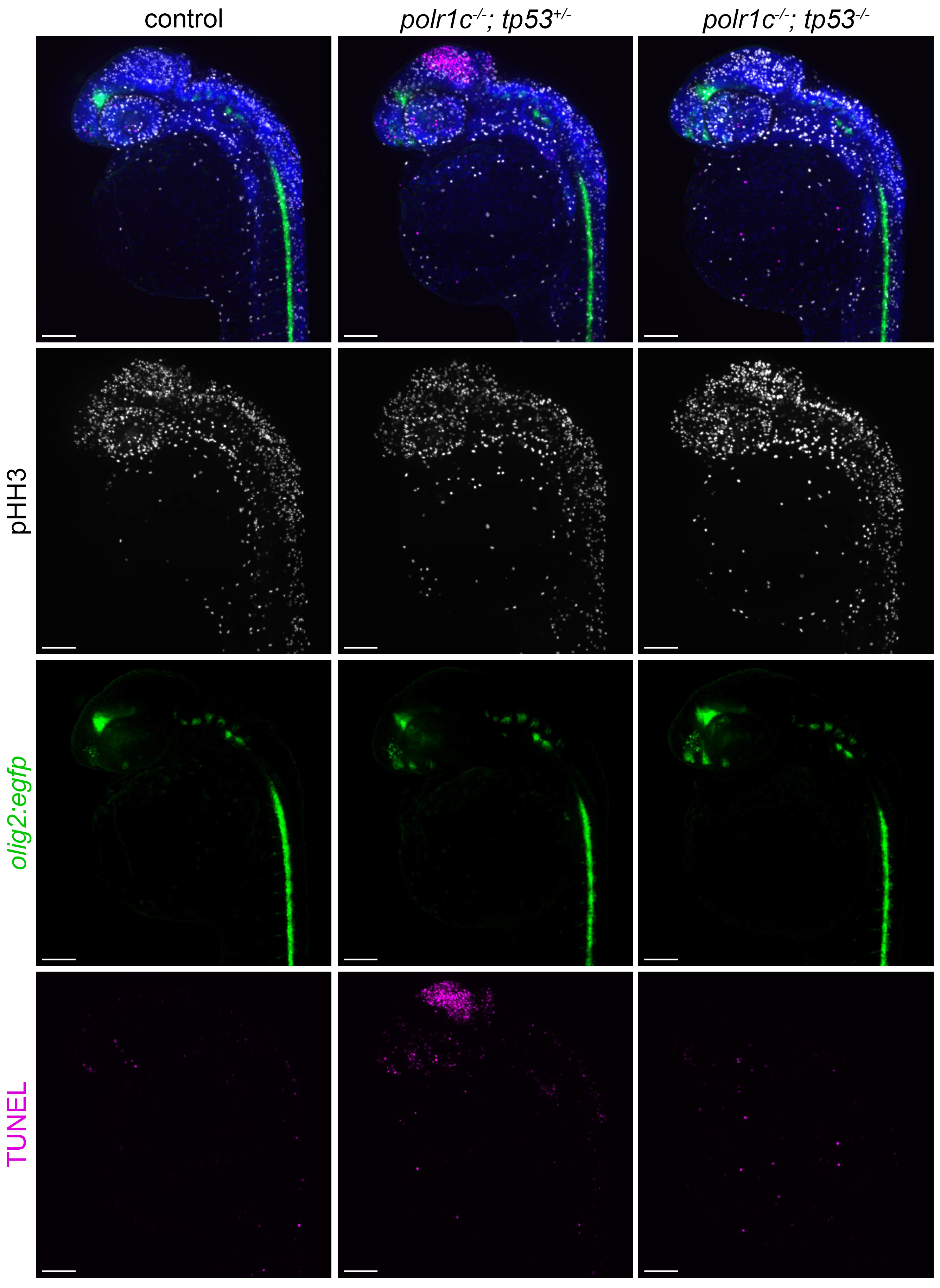

### Fig. S5

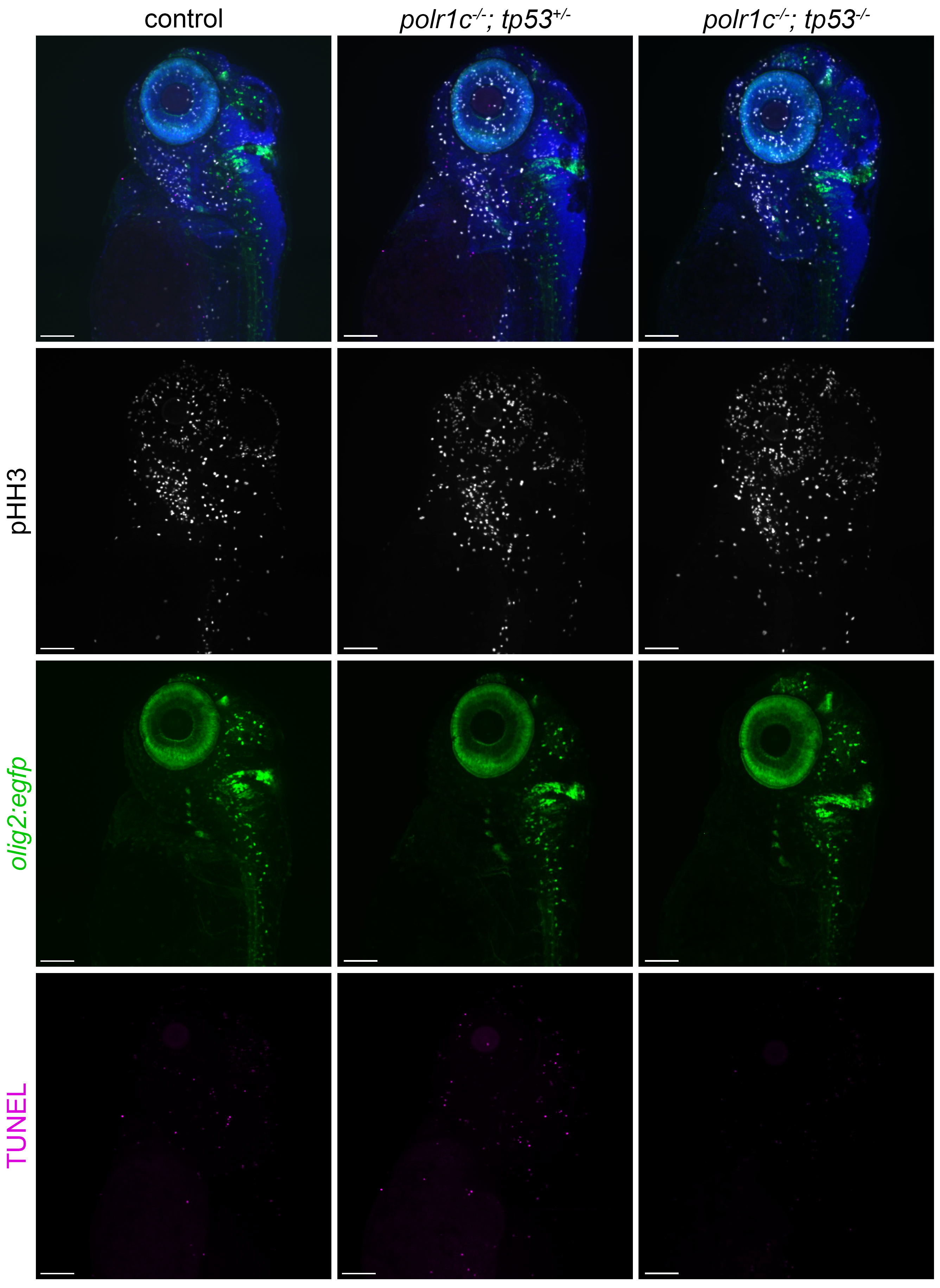

### Fig. S6

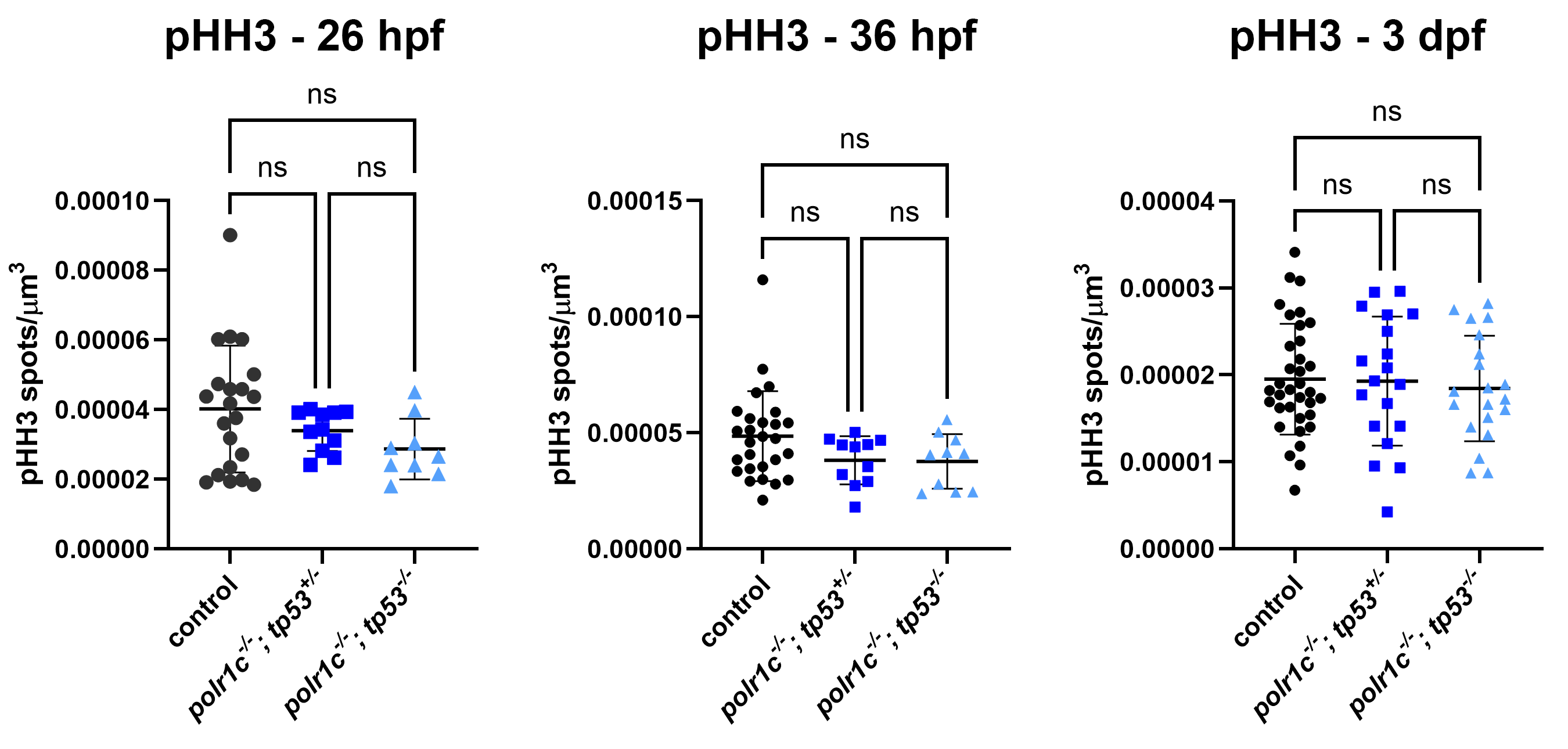
